# Geometric principles determining the morphology of oligodendrocyte precursor cells in brain white matter

**DOI:** 10.1101/2025.08.05.668685

**Authors:** Bartosz Kula, Maria Kukley

## Abstract

We used transgenic mice expressing membrane-tagged green fluorescence protein in the oligodendrocyte precursor cells (OPCs), high-resolution imaging, and detailed quantitative morphometric analysis to investigate the geometrical principles that govern structural organization of OPCs in the mouse corpus callosum. Our major findings are: (1) During the first two months of postnatal life in mice, total length of all OPCs’ processes increases via elaboration of new branches from the existing processes rather than via the appearance of new processes; (2) New branches are preferentially added to more distal sites of OPCs’ processes; (3) The processes of OPCs show stronger preferential alignment with the posterior-anterior brain axis rather than with the lateral-medial or dorsal-ventral brain axes; at the same time, the processes of OPCs show stronger preferential alignment with the lateral-medial than with the dorsal-ventral brain axis.

Our study is the first detailed comprehensive analysis of OPCs morphology comparable to those available for neurons. It helps understanding the geometrical principles that govern structural organization of OPCs. These principles are important when taking into account that OPCs receive synaptic input from neurons and are capable of synaptic integration. Arborization and structural organization of OPCs’ processes is expected to influence the travel of synaptic input from the processes (where synapses are located) to the cell soma (where synaptic inputs are integrated), in analogy to how it occurs in neurons. Hence, the integrated synaptic signal at the OPC’s cell soma which is likely to influence development and behavior of OPCs will depend on the cell morphology.

**Main Points:**

- OPC maturation increases process length through higher-order branching.
- New branches are added distally while preserving local architecture of the inner domain.
- The processes change their orientation depending on the local micro-environment.

## 1. Introduction

Oligodendrocyte precursor cells (OPCs), also known as NG2-glia, represent a distinct population of glial cells widely distributed throughout the central nervous system (CNS). One of the best-known functions of OPCs is to generate myelinating oligodendrocytes during development and also in the adult brain where OPCs persist as a self-renewing population of cells (Bergles and Richardson, 2015; van Tilborg and others, 2018). OPCs are also key players during demyelinating injury when myelinating oligodendrocytes get damaged and die, due to their capacity to proliferate, differentiate, and generate new oligodendrocytes for re-myelination of axons (Boulanger and Messier, 2014; Tepavcevic and Lubetzki, 2022). Beyond their classical role in oligodendrogenesis, accumulating evidence suggests that OPCs are functionally versatile: they are involved in synaptic signaling with neurons (Habermacher and others, 2019; Kula and others, 2019; Sun, 2024; Sun and Dietrich, 2013), participate in homeostatic and vascular regulation in support of blood-brain barrier (Akay and others, 2021; Niu and others, 2019; Seo and others, 2014), interact with extracellular matrix and potentially modulate its composition (Burnside and Bradbury, 2014; Colognato and Tzvetanova, 2011), influence adaptive and activity-dependent myelination e.g. by modifying the internode length and selection of axons to be myelinated (Fields, 2015; Mount and Monje, 2017).

The multifaceted functions of OPCs are intimately linked to their structural complexity and spatial organization within the neural tissue. Morphologically, OPCs are characterized by a small soma and a complex network of fine, highly branched processes that allow them to efficiently explore their local microenvironment (Dawson and others, 2003; Hughes and others, 2013). The dynamic nature of OPCs’ processes - constantly extending, retracting, and interacting with axons, synapses, and blood vessels (Buchanan and others, 2022; Hughes and others, 2013) - suggests that their structural organization is finely tuned to support both OPCs’ function and responsiveness to physiological and pathological changes in their environment.

However, despite the valuable existing information regarding the morphology of OPCs, our understanding of their structural organization still remains incomplete because no study has so far carried out a detailed comprehensive analysis of OPCs morphology comparable to those which are available for neurons (Halavi and others, 2012; Parekh and Ascoli, 2013). To fill this gap of knowledge, we designed a study that allowed precise tracing of all OPC processes, including the smallest ones, in a mouse line where OPCs express the *membrane-tagged* green fluorescent protein (GFP). We used high-resolution confocal imaging, carried out a very detailed manual reconstruction of the morphology of OPCs in the corpus callosum, and performed Sholl analysis, branch-order analysis, branch-angle analysis, i.e. all the morphometric techniques which were previously utilized in neurons. We focused on three age-groups: the early postnatal age at the onset of myelination (∼1,5 weeks) where there is no/very little myelin present (Chen and others, 2018; Sturrock, 1980), the juvenile age (∼3 weeks), characterized by extensive myelination in the corpus callosum reaching a peak (Chen and others, 2018; Sturrock, 1980), and the adult age (∼7 weeks) when callosal myelination is largely complete, but still continues at a slow rate (Chen and others, 2018; Sturrock, 1980). We think that the large number of cells analyzed in our study reflects the dynamic behavior of OPCs’ processes observed *in vivo*, and that we have encountered many of the OPCs’ dynamic states.

We present here the results of our work. We think that our quantitative morphometric analysis and systematic characterization of the morphological features of OPCs helps understanding the geometric principles that govern structural organization of OPCs. Knowledge of these principles is important due to OPC’s ability to receive and integrate synaptic input from neurons (Kula and others, 2019; Sun, 2024; Sun and Dietrich, 2013), which drive their fate (Chen and others, 2018; Kula and others, 2019; Nagy and others, 2017). Arborization and structural organization of OPCs’ processes will influence the travel of the synaptic input from the processes (where synapses are located) to the OPCs cell soma (where synaptic inputs are integrated), in analogy to how it occurs in neurons (Grienberger and others, 2015; Olcese and others, 2013; Stuart and Spruston, 2015). Hence, the synaptic signal that arrives at the OPC’s cell soma, and is subject to the integration, will depend on the cell morphology.

## 2. Materials and Methods

### 2.1. Ethics statement

All experiments were performed in accordance with current European Union guidelines and approved by the local government authorities for Animal Care and Use (Regierungspraesidium Tübingen, State of Baden-Wuerttemberg, Germany). All efforts were made to minimize the suffering of the animals.

### 2.2. Animals

In all experiments we used the offspring mice from the cross-breeding of two mouse lines: the B6.129(Cg)-Gt(ROSA)26Sortm4(ACTB-tdTomato,-EGFP)Luo/J (ROSAmT/mG) and the B6.Cg-Tg(Cspg4-cre/Esr1*)BAkik/J (NG2CreERTM). The breeding pairs were initially obtained from the Jackson Laboratory (stocks: #007676 and #008538, respectively), and subsequently maintained at the Animal Facility of the Hertie Institute of the University of Tübingen. All animals were housed under 12/12 hour light/dark conditions with water and food available *ad libitum*.

### 2.3 Tamoxifen injection

The activity of Cre-recombinase in the postnatal ROSAmT/mG:NG2-CreERTM double-transgenic mice was induced by intraperitoneal injection of 4-hydroxytamoxifen (4-OHT, Sigma). A 10 mg/ml stock solution was prepared by dissolving 4-OHT in 19:1 autoclaved vegetable oil:ethanol. Mice were injected with 1 mg of 4-OHT per gram of body weight intraperitoneally at the age of P8-11, P9-22 and P50-53. Each animal received one injection of tamoxifen. P19-22 and P50-53 animals were anesthetized with a low dose of Isoflurane for the procedure. Control animals were injected with the same volume of vehicle, i.e. 19:1 autoclaved vegetable oil:ethanol. We performed anti-GFP staining also in the control animals, but have not detected any signal. Therefore, these animals were not further analyzed.

### 2.4. Preparation of brain slices

Three days after the 4-OHT injection, mice were anesthetized with a mixture of Isoflurane (3% v/v) in 100% oxygen, and decapitated. Whole brains were removed and 300 μm coronal brain sections were prepared in ice-cold carbogenated (95% O2, 5% CO2) N-methyl-D-glucamine (NMDG) solution containing (in mM): 135 NMDG, 1 KCl, 1.2 KH2PO4, 20 choline bicarbonate, 10 glucose,1.5 MgCl2 and 0.5 CaCl2 (pH 7.4, 310 mOsm) using Leica VT 1200S vibratome. Subsequently, the slices were transferred into 4% paraformaldehyde (PFA) in 0.01M phosphate buffered saline (PBS) and fixed for 20-24 hours at 4°C. Afterwards slices were washed in PBS, embedded into 5% agar in PBS and re-sectioned to 70 μm at Thermo Fisher Scientific HM 650V microtome. Slices were washed with 0.1M Tris-buffered saline (TBS, pH 7.6) three times. Afterwards, a blocking solution containing 0.1M TBS, 10% fraction V albumin (Roth) and 1% Triton-X (Roth) was applied for 1 hour at 37°C. Afterwards slices were incubated for 48 hours with primary antibodies (1:500) in 0.1M TBS, 10% albumin V and 0.5% Triton-X at 4°C. We used the following primary antibodies: guinea pig or rabbit anti-NG2 antibodies (gifts from Dr William Stallcup, Burnham Institute, La Jolla, CA, USA); rabbit anti-GFP antibody (Invitrogen) or chicken anti-GFP antibody (Abcam). After a short wash in TBS slices were incubated with secondary antibodies (1:1000) in TBS, 10% albumin and 0.5% Triton-X for 48 hours at 4°C. We used the following secondary antibodies: anti-chicken Alexa Fluor 488 (Invitrogen), or anti-rabbit Alexa Fluor488 (Invitrogen), or anti-chicken fluorescein isothiocyanate (FITC, Dianova) to detect GFP; anti-rabbit Alexa Fluor633 (Invitrogen), or anti-guinea pig Alexa Fluor633 (Invitrogen), or anti-rabbit Rhodamine Red-X (RRX, Dianova) to detect NG2. Cell nuclei were labelled by incubating the slices for 20 min with 0.2 μg of 4’6-diamidino-2-phenylindole (DAPI) in TBS. After washing with TBS, the slices were left to air-dry for 10 min. Subsequently, they were mounted on glass slices in Vectashield Antifade Mounting Medium (Vector Labs), and sealed with nail-polish. Preparations were stored in the dark at 4°C until confocal microscope imaging.

### 2.5. Image acquisition

Immunostained sections were imaged at Zeiss LSM 710 Meta Confocal microscope equipped with 63x Plan Apochromat oil immersive objective (NA=1.4). Pinhole size was set to 50 μm (i.e. approximately 1 Airy unit for 488 nm wavelength). Whole cells were scanned using a zoom factor of 1.1 – 1.5, pixel dwell ranging 9.7-12.5 μs, and x,y pixel size of 0.089. The z-step between images within a stack was set to 0.37μm, resulting in a 50% overlap between the optical sections (0.74 μm/section). Frame size was adjusted to keep identical pixel size at every zoom level. All images were acquired with 16 bit color depth and 4x averaging. The following laser excitation lines and emission detection ranges were used: for Alexa Fluor 488/FITC, λ_ex_ = 488 nm, λ_em_ = 494-553 nm; for Alexa Fluor 568/RRX, λ_ex_ = 561nm, λ_em_ = 562-631nm; for Alexa Fluor 633 λ_ex_ = 633 nm, λ_em_ = 641-729; for DAPI, λ_ex_ = 410 nm, λ_em_ = 416-474. Laser power of 1.5 – 3.0% was used for 488 nm, 561 nm and 633 nm lasers. Laser power of 0.3-0.5% was used for the 405nm laser. Laser power, gain and offset were adjusted for the best possible signal to background ratio. All acquired stacks of images were saved in Zeiss .lsm file format which preserves the meta-data information.

### 2.6. Cell tracing

Previously acquired confocal image stacks in .lsm format were uploaded into Neurolucida (MBF Bioscience, USA). The voxel sizes were loaded automatically from the .lsm files by the software. Single pixel-thick cell reconstructions were created by tracing manually through the middle of the processes. Brightness/contrast and gamma were adjusted during tracing to keep similar intensity profiles at all traced slices/sections. Every tubular or conical protrusion originating from the soma was classified as a process, regardless of length or branching.

### 2.7. Branch order and the positioning of process elements

In our study, a branch is defined as a part of a process located between either: (1) the origin point (root) and a branching point (node); or (2) between two branching points; or (3) between a branching point and a process ending. Centrifugal ordering, which assigns numbers in an ascending order, starting from process origin, was used to mark the position of new branches along the process. Briefly: a branch starting at the origin of a process (root) is assigned the order=1. If the branch terminates with a branching point, this branching point is also assigned the order=1. All new branches starting at the branching point are assigned the order=+1. Endings are ordered in the same way and receive the branching order of the branch which they terminate. This numeration continues until all branches, branching points, and endings have an order assigned. As an example: if a branch originates from a branching point of the order=3, then the branch is of order=4, and its ending is also of order=4.

### 2.8. Sholl analysis

Sholl analysis was performed in the Neurolucida Explorer. A set of concentric shells spaced by 2.5 μm and anchored at the centroid of the cell soma was placed over each OPC. The shells partition the brain parenchyma into volume compartments separated by the shells. The intersections between the shells and the OPCs’ processes indicate the number of times those processes cross each shell at a given radius from the center of the cell soma. These intersections represent a measure of the complexity and extent of the arbors of OPCs’ processes. We quantified the number of the branching points, endings, and branches, as well as the length of those branches, contained within individual compartments, and analyzed them in relation to increasing distance from the center of cell soma. Note, that the plots on the relevant figures indicate only the upper radius of the shell range.

### 2.9. Nearest Neighbor Analysis for Branching Points

To examine the number and distances between neighboring branching points, we first calculated the Euclidean distances between all branching points of a cell, irrespective of their position, process, or branch order. We then performed two complementary analyses:

First, using principles similar to those in the Sholl analysis (Section 2.8), we excluded all neighbors located beyond a specified cutoff distance from each selected point. Cutoff values ranged from 1□µm to 5□µm, in 1□µm increments. For each branching point, we determined the number of neighbors within each cutoff distance and plotted this number against either the point’s distance from the soma centroid (binned and averaged every 2.5□µm, up to 40□µm) or its branch order.

Second, we calculated the Euclidean distances between each selected branching point and its five closest neighbors. These distances were then plotted against the point’s distance from the soma centroid or its branch order, as described above.

Results are presented as mean values of averaged neighbor counts and distances for each experimental group.

### 2.10. Branch direction in space and alignment with brain axes

To study changes in the direction of branches, we converted the Cartesian coordinate system used by Neurolucida into a spherical coordinate system. In the spherical coordinate system, position of a point is described by three values: (1) the radial distance r, which is the distance towards beginning of the coordinate system; (2) the azimuth angle φ defined for x,y axes, and (3) the polar angle θ corresponding to the z axis.

For coronal slices, used in our study, the x and y axes are aligned with the lateral-medial (L-M) and dorsal-ventral (D-V) anatomical axes, respectively. The z axis aligns with the anterior-posterior axis (A-P).

The azimuth angle φ takes values between 0° and 360°. Therefore, to measure the alignment of branches with L-M and D-V axes, φ quadrants were converted to 0°-90° where angles approaching 0° align with L-M axis while angles approaching 90° align with D-V axis.

The polar angle θ takes values from -90° to +90°, but we converted them to 0°-90°, where angles approaching 90° align with the A-P axis while angles approaching 0° completely miss-align with the A-P axis.

Due to the tortuosity of branches, all angular measurements were performed on vectors spanning over the origin of the branch (root or branching point) and its terminus (another branching point or ending). The lengths reported are the real lengths of the branches, not vector lengths. For graphing and statistical analyses all measurements were grouped into 15° bins, i.e. 6 bins in total. The radial distance was ignored and φ and θ angles were analyzed separately.

### 2.11. Change in branch direction after a branching point

To measure changes in direction, all branches were approximated as vectors spanning over their origins and terminations. For each pair of a “mother” branch (terminating at a branching point) and its “daughter” branch (originating from that branching point) the planar angle was measured as an angle between the “daughter” vector and a linear extension of the “mother” vector. In the measurement, the “mother” and the “daughter” vectors are assumed to share a unique 2D plane. If the “daughter” branch does not change direction, the angle is 0°; if the “daughter” branch is parallel to its “mother” branch but is facing backwards, the angle is 180° (which is the largest possible angle).

### 2.12. Statistics

Statistical analysis was performed in Graph Pad Prism 9.3.1. All datasets were tested for homoscedasticity and normality. If the datasets met the assumptions of normality and homogeneity of variances, one-way ANOVA followed by post hoc Holm-Šídák’s test was used. If the datasets did not meet the assumption of equality of variances (regardless of normality), Brown-Forsythe and Welch ANOVA followed by post hoc Dunnet’s T3 test were used. If the datasets did not meet the assumptions of normality, but had equal variances, Kruskal-Wallis test followed by post hoc Dunn’s test were used. If the data were plotted against a continuous factor (such as binned distance in Section 2.9) and met the assumptions of normality and homogeneity of variances, a two-way ANOVA followed by Holm-Šídák’s post hoc test was used. For all statistical comparisons, the significance level was set at p < 0.05. Statistically significant differences are indicated on the figures by markings: ∗ represents p ≤ 0.05, ∗∗ represent p ≤ 0.01, and ∗∗∗ represent p ≤ 0.001. The exact p values are given in the text. If the data is normally distributed, the data in the text or figures is reported as scatter plots and mean ± standard error of the mean (SEM) or, if not normally distributed as a boxplot containing median and 10th, 25th, 75th, 90th or 25th, 75th percentiles.

The datasets were checked for outliers by Prism’s ROUT method at Q=5%.

All data acquisition was randomized. Throughout the study we made all efforts to avoid pseudo replication by restricting the maximum number of cells acquired from a single animal to three. The exact number of cells and animals used in each experiment is given in the figure legends.

### 2.13. Power and sample size calculations

The sample sizes for all of the statistical comparisons were determined based on the mean values and pooled standard deviations from preliminary tracings of 8-10 cells per group, α = 0.05, β = 0.8, corrected for the number of pairwise comparisons (k), based on the following equations:

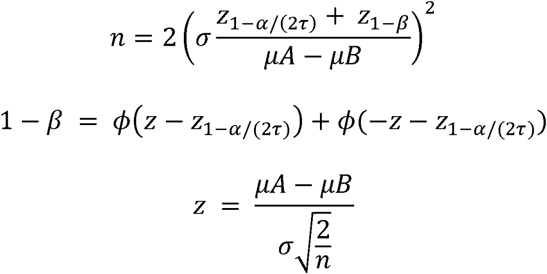

where n is the sample size; σ is standard deviation; is standard Normal distribution function; α is Type I error; is the number of pairwise comparisons; β is Type II error. During the calculations, the normality of residuals and equality of variances were assumed a priori.

The calculations were performed in an online calculator available at: http://powerandsamplesize.com/Calculators/Compare-k-Means/1-Way-ANOVA-Pairwise

## 3. Results

### 3.1. Choice of animals for structural analysis of callosal OPCs

The goal of the present study was to perform a detailed reconstruction of the morphology of OPCs in the mouse corpus callosum. Therefore, we sought for an approach that allowed precise tracing of all OPC processes, including the smallest ones. To achieve this, we used a mouse line in which OPCs express the membrane-tagged green fluorescent protein (GFP). These animals were generated by crossing NG2CreER^TM^ mice, where inducible Cre-recombinase is expressed under the control of NG2 promoter (Zhu and others, 2011; Zhu and others, 2012) and ROSA^mT/mG^ reporter mice, where two *lox P* sites flank a STOP codon in front of the GFP cassette (Zhu and others, 2011; Zhu and others, 2012).

Expression of GFP was activated by two injections of 4-OH tamoxifen (1 mg/g body weight) in mice of the three age-groups, and animals were sacrificed 3 days after the injection (Figure 1A). The relatively low dose of 4-OH tamoxifen was chosen to achieve sparse GFP-labelling of OPCs; this was important for clear separation of individual OPCs from each other, avoiding the intermingled processes between neighboring cells that could pose challenge during cell tracing (Figure 1B-D). Notably, 3 days after 4-OH tamoxifen injection not only OPCs but also few oligodendrocytes (presumably differentiated from the GFP-labelled OPCs) expressed GFP (Figure 1B-D,H). Therefore, we performed immunolabelling for NG2, which is a marker of OPCs, and included into the analysis only the NG2^+^GFP^+^ cells (Figure 1E-H). As NG2 also labels pericytes which are the components of blood vessels, we took care that NG2^+^GFP^+^ cells targeted for tracing and reconstruction were located away from blood vessels (Fig. 1F, H).

**Figure 1:**
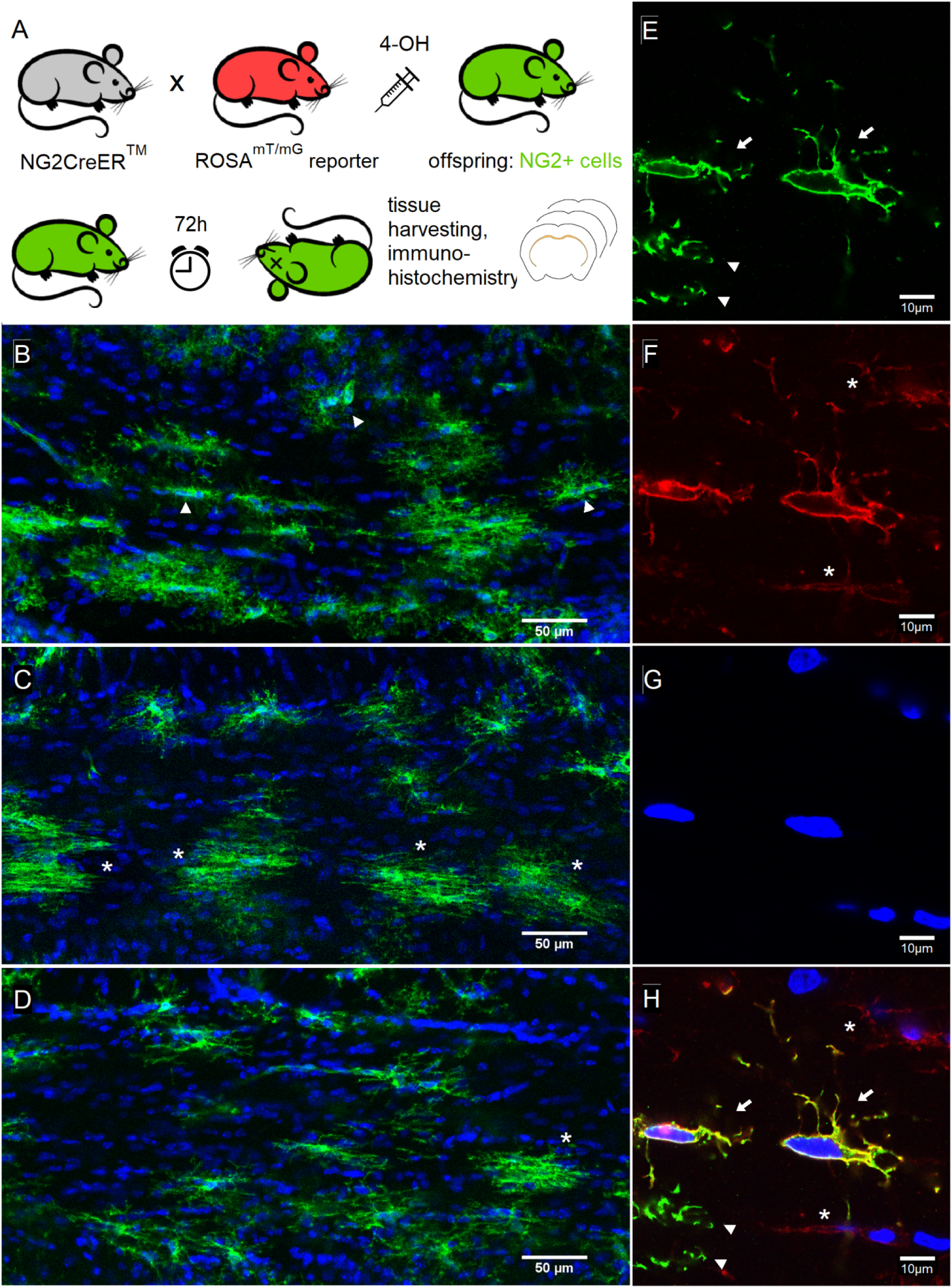
Tamoxifen injections for sparsely labelling of callosal OPCs. **(A):** Experimental design: The progeny of NG2CreER^TM^ and ROSA^mT/mG^ mouse lines received tamoxifen and were sacrificed 72h later. **(B) - (D):** 10x magnified confocal images of the corpus callosum stained for GFP (green) and DAPI (blue) at P10 **(B)**, P20 **(C)** and P50 **(D)**. **(E) - (H):** Immunohistochemical verification of GFP^+^ cells used for tracing: GFP **(E)**, NG2 **(F)**, DAPI **(G)** and all three channels merged **(H)**. The GFP^+^ NG2^+^ cells are marked with arrows, GFP^+^ NG2^-^ cells marked with arrowheads, GFP^-^ NG2^+^ cells marked with asterisks. **(I)**: Maximum intensity projection of a confocal stack containing a GFP^+^ NG2^+^ cell used for tracing. Neighboring GFP^+^ cells were removed for clarity. **(J)**: Tracing of the cell shown in (I).

We reconstructed all cells in Neurolucida as single-pixel skeletons (Figure 2A-E). This approach allowed us to observe that OPCs show diverse morphology in animals of all three investigated ages (Figure 2A-E). We analyzed the number and architecture of the OPC processes, including length, branching, and orientation. We also looked in detail at the branching patterns of the OPCs’ processes trying to define the geometrical principles guiding the arborization. We compared the data between the three age-groups of mice in order to find out how structural changes of OPCs may correlate with maturation of the corpus callosum and myelination.

**Figure 2:**
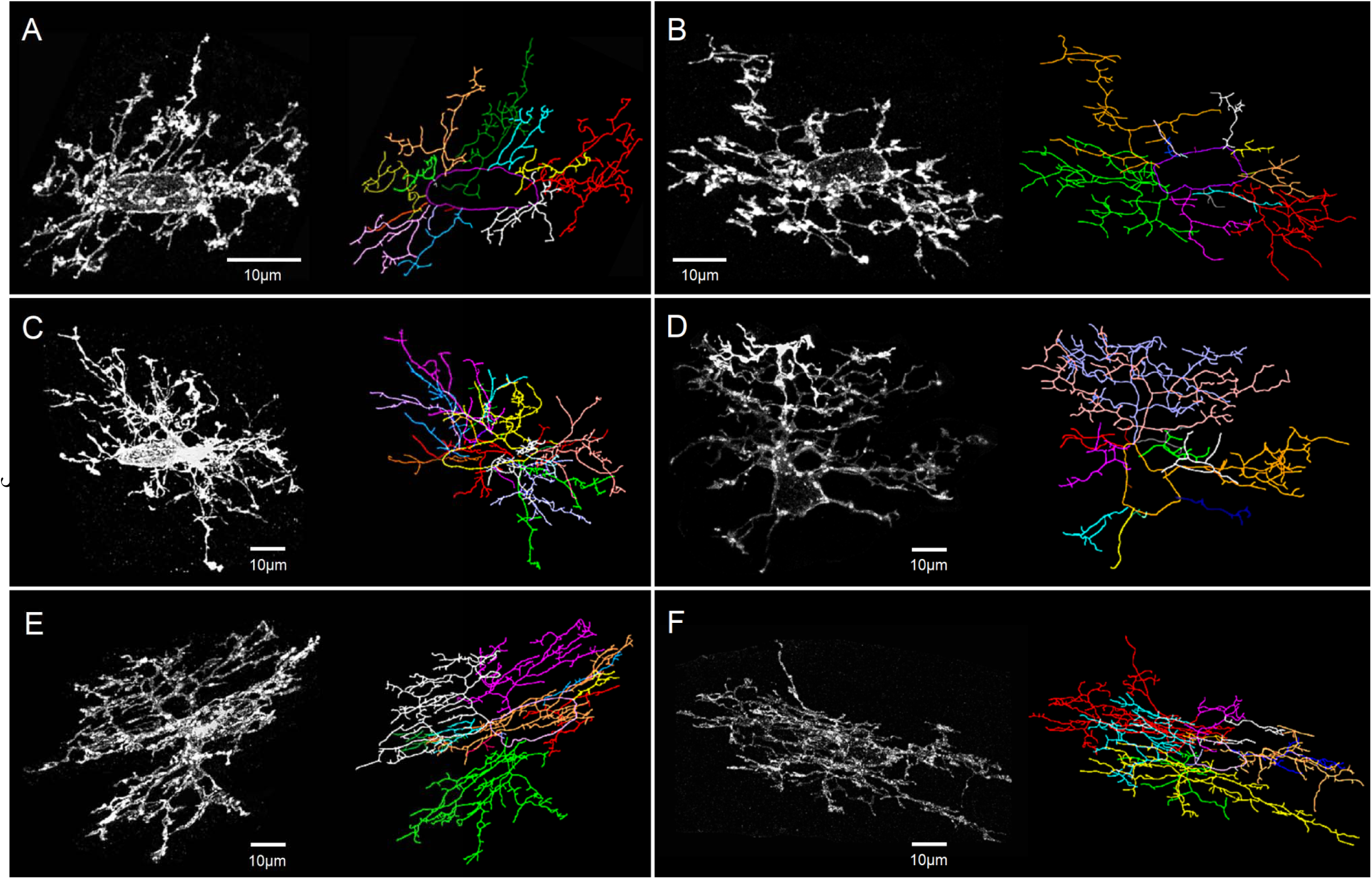
Callosal OPCs display greatly diverse morphologies at all investigated ages. **(A):** Maximum intensity projection of a P10 OPC (left) and the corresponding tracing (right). A young, very flat OPC possessing clearly distinct processes of mostly similar length. Each process is labeled with a different color. Soma outline is in purple. **(B):** Maximum intensity projection of another P10 OPC and the corresponding tracing. In contrast to (A), this young OPC has 3 large, massive processes (green, orange, red) and multiple much shorter ones. **(C):** Maximum intensity projection of a P20 OPC and the corresponding tracing. Note a remarkable difference in the length of processes when compared with both P10 cells. This cell has multiple processes of comparable length which frequently intermingle. Soma is outlined in yellow. **(D):** Maximum intensity projection of another P20 OPC and the corresponding tracing. Note irregular shape of the soma and the lack of bipolar morphology with an unusual asymmetry of process distribution. This cell has 3 large processes (indicated in violet, pink, and orange) which are all localized at the same (top-right) side of the soma. Only short processes originate from the left-bottom side of the soma. Soma is outlined in orange. **(E):** Maximum intensity projection of a P50 OPC and the corresponding tracing. The cell does not have bipolar morphology (although soma is oval and regular), and seems to be selective towards the direction of its processes, as in (D). In addition, clear gaps in the process coverage of the surrounding brain parenchyma are visible. Soma is outlined in pink. **(F):** Maximum intensity projection of another P50 OPC and the corresponding tracing. This cell has processes strongly aligned with the callosal axons. Soma is outlined in pink.

### 3.2. Number of processes in OPCs does not change as animals mature but the total length of all processes increases

We first studied the general morphology of OPCs in the three age-groups of mice by counting the number and the length of their processes. We defined a process as a structure originating at the cell soma; it could be very short and have no branches, or it could be very long and extensively branched. We defined a branch as a part of a process located between two branching points, or between a branching point and an ending point (Figure 3A, B).

**Figure 3:**
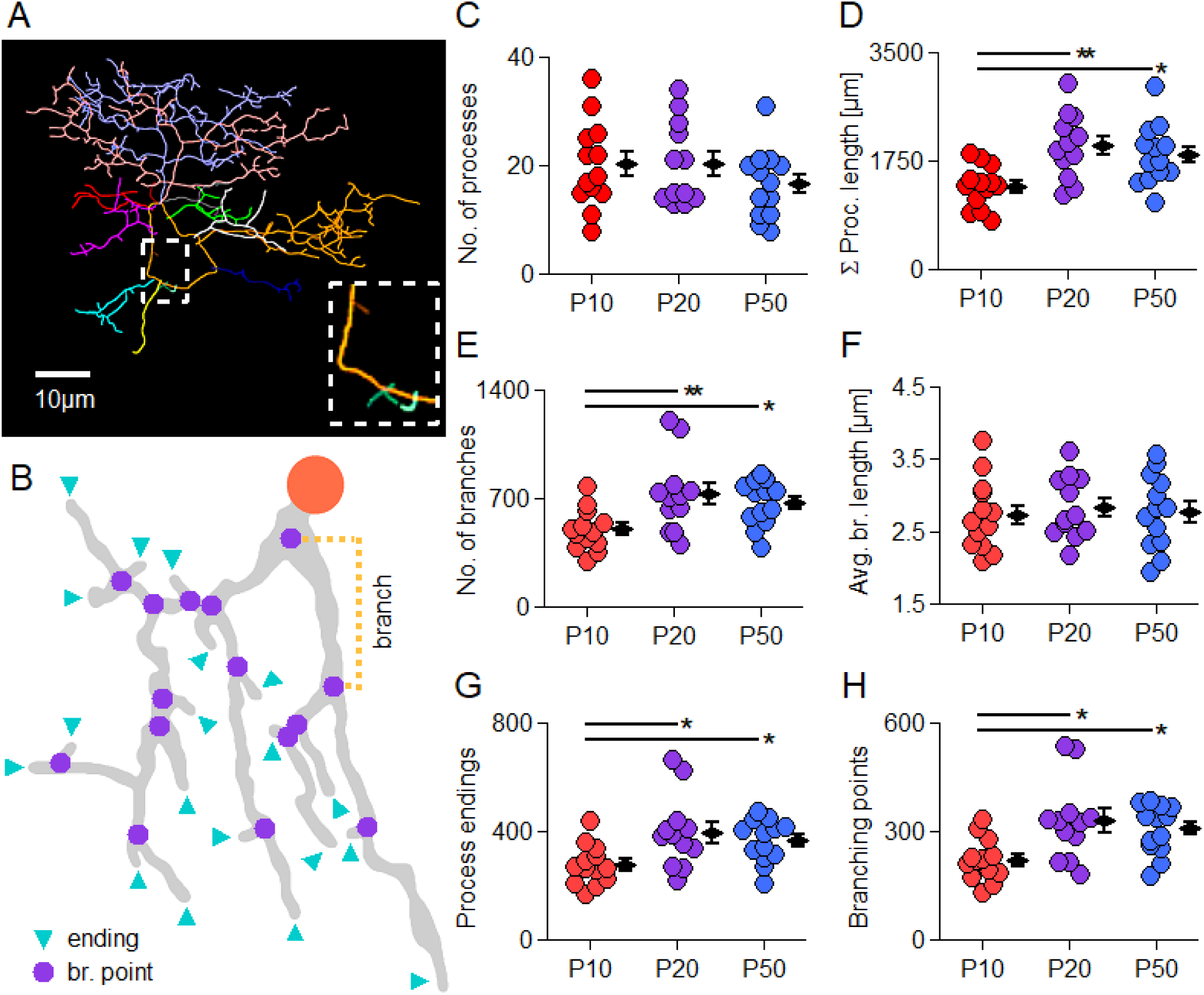
Number of processes in OPCs does not change as animals mature but the total length of all processes increases. **(A):** An example of an OPC without a bipolar morphology. Each process is labeled with a different color. Note the large diversity in the length of processes and branches, and the uneven space filling by the processes. The inset shows three tiny filopodia processes; larger processes were removed from the inset for clarity. **(B):** Schematic drawing of a single process. The origin at the soma is represented by a large red circle. Small violet circles represent branching points of the process. Blue triangles point towards process endings. A branch is a part of the process between a branching point and another branching point or an ending. An example of a branch is marked by an orange semi-box. **(C):** Comparison of the total number of processes per cell. All processes in an individual cell were counted regardless of their length or branching. Each circle represents the count in one cell. Black diamonds represent group average ± SEM. Statistically significant differences are highlighted by horizontal bars, *=p<0.05, **=p<0.01. P10 group: n=13 in 7 animals; P20 group: n=12 in 8 animals; P50 group: n=13 in 8 animals. The same labeling, number of cells (n), and the number of animals apply to the other graphs within this figure. **(D):** Total length of all processes within each cell, compared between the age-groups. **(E):** Total number of branches within each cell, compared between the age-groups. Branches are counted regardless of the process length or branching. **(F):** Total number of branching points in each cell, compared between the age-groups. **(G):** Total number of endings in each cell, compared between the age-groups. **(H):** Average length of a single branch per cell, compared between the age-groups.

We found that the total number of processes did not differ significantly between the age-groups: 20.15 ± 2.19 at P10; 20.33 ± 2.17 at P20; and 16.62 ± 1.76 at P50 (p=0.357) (Figure 3C). The variability of this parameter, estimated as the coefficient of variation (CV), was high but nearly identical between the groups: 8-36 processes, CV=0.40 at P10; 13-34 processes, CV=0.38 at P20; 8-31 processes, CV=0.39 at P50. These findings compare well with a previous study showing that callosal OPCs may have 8-30 process-trees, although that study focused on OPCs in older (3 to 11 months old) animals (Dawson and others, 2003).

Next, we investigated the total length of all processes in OPCs which was calculated as a sum of the lengths of all branches. We found ∼30% increase at P20 and P50 mice compared to P10 animals: 1340.44 ± 92.34 µm at P10; 2002.62 ± 152.94 µm at P20; 1851.85 ± 132.65 µm at P50 (P10 vs P20 p=0.002; P10 vs P50 p=0.013; P20 vs P50 p=0.410, Holm-Šídák’s test) (Figure 3D).

### 3.3. Higher branching underlies larger total process length in more mature OPCs

Each process in an OPC may be built by a single branch or by multiple branches. Therefore, the observed increase in the total length of all processes in OPCs upon animal maturation (Figure 3D) may be caused by elongation of individual branches without change in their number, by increase in the number of branches without change of their length, or by alterations of both parameters. To distinguish between those possibilities, we analyzed the branch length, the number of branches, as well as the number of branching points and the endings.

We found that the number of branches was ∼30% higher in OPCs from P20 and P50 animals compared to P10 mice (Figure 3E): 498.54 ± 36.22 at P10; 725.67 ± 70.36 at P20; 674.69 ± 40.54 at P50 (P10 vs P20 p=0.009; P10 vs P50 p=0.033; P20 vs P50 p=0.48, Holm-Šídák’s test). Similar changes were observed in the number of branching points (Figure 3F): 221.00 ± 16.13 at P10; 328.92 ± 31.70 at P20; 307.46 ± 18.88 at P50 (P10 vs P20 p=0.006; P10 vs P50 p=0.020; P20,P50 p=0.510, Holm-Šídák’s test), and in the number of ending points (Figure 3G): 277.54 ± 20.30 at P10; 396.67 ± 38.75 at P20; 367.23 ± 21.72 at P50 (P10 vs P20 p=0.013; P10 vs P50 p=0.050; P20 vs P50 p=0.46, Holm-Šídák’s test). The identical increase in all three parameters suggests that addition of new branches rather than elongation of the existing ones is responsible for the observed increase in the total length of OPCs processes upon animal maturation. Indeed, when we analyzed the mean length of individual branches, we found that it remained almost identical throughout the development (Figure 3H): 2.73 ± 0.14 µm at P10; 2.83 ± 0.12 µm at P20; 2.77 ± 0.14 µm at P50 (one-way ANOVA, p=0.867).

Taken together, our data show that a larger total length of OPC processes in the more mature animals is caused by a stronger branching of the OPC processes.

### 3.4. In more mature animals, new branches are preferentially added to more distal sites of OPC processes

We were wondering whether stronger branching of OPC processes in more mature animals implies that new branches are added equally throughout the whole length of processes or rather at a certain preferential location. To test this, we used Sholl analysis which is a well-established technique to compare the branching and the distribution of cellular processes in space (Sholl, 1953). For this analysis we drew a set of concentric shells spanning over the whole volume occupied by a given OPC, with the center of each shell located at the centroid of the cell soma (Figure 4A). We set the spacing between the shells to 2.5 μm (Figure 4A) to roughly match the average branch length. To evaluate how the structure of processes changes with increasing distance from the cell soma, we analyzed the number of intersections between the process skeletons (single-pixel approximations preserving lengths but ignoring local thickness) and the individual shells (Figure 4A). To evaluate how the number of the branching points and the endings, as well as the length of individual branches, changes depending on the distance from the cell soma, we analyzed those parameters within the volume compartments located between two neighboring shells (Figure 4B).

**Figure 4:**
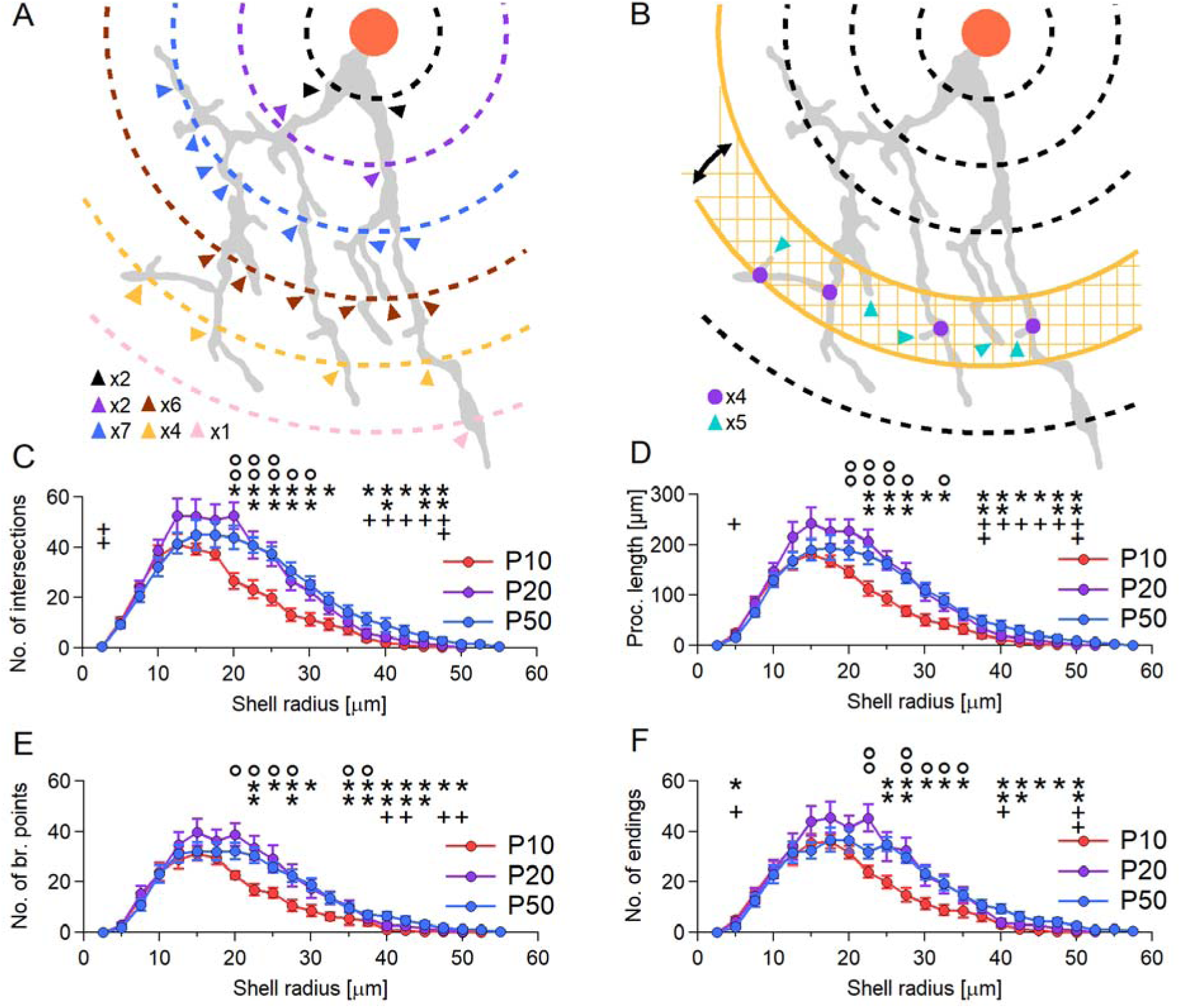
During animal maturation, OPCs preserve the process density within the inner part of the cell domain but increase the density in its outer sites. **(A)**: Example of Sholl analysis: a set of concentric shells (represented in 2D as circles), spaced every 2.5 μm, is positioned over the centroid of the cell soma. Then, the number of intersections between the process and each of the shells is counted and plotted against the shell’s distance from the soma (radius). In this example, each consecutive shell has a different color. Arrowheads point to the intersections with the specific shell. Bottom left: a summary of the number of intersections of the process with each shell. **(B):** The shells partition the brain parenchyma into volumetric compartments and allow for measurement of the structure lengths and/or the number of sub-structural elements contained within the shells. In this example, arrows show a shell spun between 2 shells (in orange) singled for the analysis. The length of the process, the number of branching points or endings contained in the checked area (volume) is measured and plotted against the upper radius of the shell range (for example the point of 30 μm on the graph corresponds to the spun between 27.5 μm and 30 μm). **(C):** Comparison of the number of intersections with the shells spaced every 2.5 μm. All points on the graph represent the group averages ± SEM: P10 in red, P20 in green, P50 in blue. Significant differences are marked with ° for P10 vs P20 comparisons; * for P10 vs P50 comparisons; ^+^ for P20 vs P50 comparisons. °,*,^+^ = p<0.05; °°,**,^++^ = p<0.01. P10 group: n=13 in 7 animals; P20 group: n=12 in 8 animals; P50 group: n=13 in 8 animals. **(D):** Comparison of the length of processes within shells spaced every 2.5 μm. Data is plotted against the upper radius of the shell range (for example the point of 30 μm on the graph corresponds to the shell spun between 27.5 μm and 30 μm). The labeling, n numbers and plotting radius are identical to (C). **(E):** The same as (D) for the number of branching points within shells. The labeling, n numbers and plotting radius are identical to (C). **(F):** The same as (D) for the number of endings within shells. The labeling, n numbers and plotting radius are identical to (C).

First, we looked into the number of intersections plotted against the distance from the cell soma center (Figure 4C). We found that up to the distance of ∼17.5 μm, the number of intersections was similar between the age groups (Figure 4C; one-way ANOVA p=0.111 to 0.808). Furthermore, in all groups the number of intersections reached the peak at the distance of ∼15 ± 2.5 μm and declined thereafter (Figure 4C; 39.154 ± 2.275 at P10; 52.167 ± 6.882 at P20; 44.769 ± 5.876 at P50; one-way ANOVA p=0.268). The first significant differences between groups were visible at the distance of ∼20 μm: in the P10 group, the number of intersections declined rapidly, while in P20 and P50 groups it remained comparable to the peak (Figure 4C; 26.539 ± 3.010 at P10; 52.500 ± 5.373 at P20; 43.615 ± 4.509 at P50; P10 vs P20, p=0.000543; P10 vs P50, p=0.0160; P20 vs P50, p=0.161]. At distances above 20 µm, all three age-groups showed a fast progressing decline (Figure 4C). The difference between the P10 and the P50 groups was statistically significant almost throughout the whole distance range above 20 µm, up to the last detected intersections at ∼55 μm (Figure 4C). On the other hand, the difference between the P10 and P20 groups was statistically significant only throughout the distance of ∼20-30 μm. Afterwards the P20 distribution remained statistically similar to P10, but became different from P50 (Figure 4C), suggesting that OPCs in P50 animals extend their processes the furthest among the age groups.

Next, we analyzed the total length of all processes, compartmentalized by Sholl shells at various distances from the cell soma, as well as the number of branching points and endings in those compartments (Figure 4B). We found that those distributions (Figure 4D, E, F) closely followed the distribution of intersections (Figure 4C): peaked at the distance of ∼15 ± 2.5 μm for the length (Figure 4D) and the number of branching points (Figure 4E), and at 17.5 ± 2.5 μm for the number of endings (Figure 4F). The total lengths of processes at peak were: 180.854 ± 11.897 at P10; 241.158 ± 32.382 at P20 (but note the high variability); 189.031 ± 22.039 at P50 (Figure 4D; P10 vs P20 p=0.209; P10 vs P50 p=0.801; P20 vs P50 p=0.229). The numbers of branching points at peak were: 30.923 ± 2.908 at P10; 39.583 ± 5.226 at P20; 31.923 ± 3.529 at P50 (Figure 4E; P10 vs P20 p=0.345; P10 vs P50 p=0.857; P20 vs P50 p=0.345). The number of endings at peak: 35.308 ± 3.461 at P10; 43.9167 ± 5.796 at P20; 32.231 ± 3.296 at P50 (Figure 4F; P10 vs P20 p=0.301; P10 vs P50 p=0.607; P20 vs P50 p=0.174). The overall shapes of those three distributions were comparable to the corresponding distributions of the numbers of intersections (compare Figure 4D, E, and F to Figure 4C). Furthermore, the differences between the age-groups in the distributions of the process lengths, numbers of branching points and number of endings tested as statistically significant throughout the same ranges of distances as for the corresponding distributions of the intersections.

Taken together, our findings indicate that stronger branching of OPC processes in more mature animals is caused by preferential incorporation of new branches at the distal sites of the processes, i.e. at locations of more than 20 µm from the cell soma.

### 3.5. Larger number of higher-order branches underlies stronger branching of OPC processes in more mature animals

Although Sholl analysis is a valuable tool for studying the distribution and density of cellular processes in space, it does not provide a structured view of the branching pattern and does not allow for the quantification of branches at different hierarchical levels. To fill this gap and to look at the structural organization of OPC processes in further detail, we performed the Branch Order Analysis (Figure 5A, B; see Materials and Methods for details). The assumptions of this analysis are almost the opposite to those of the Sholl analysis: the location in space is now ignored, and only the hierarchical level of the position on the process is treated as relevant.

**Figure 5:**
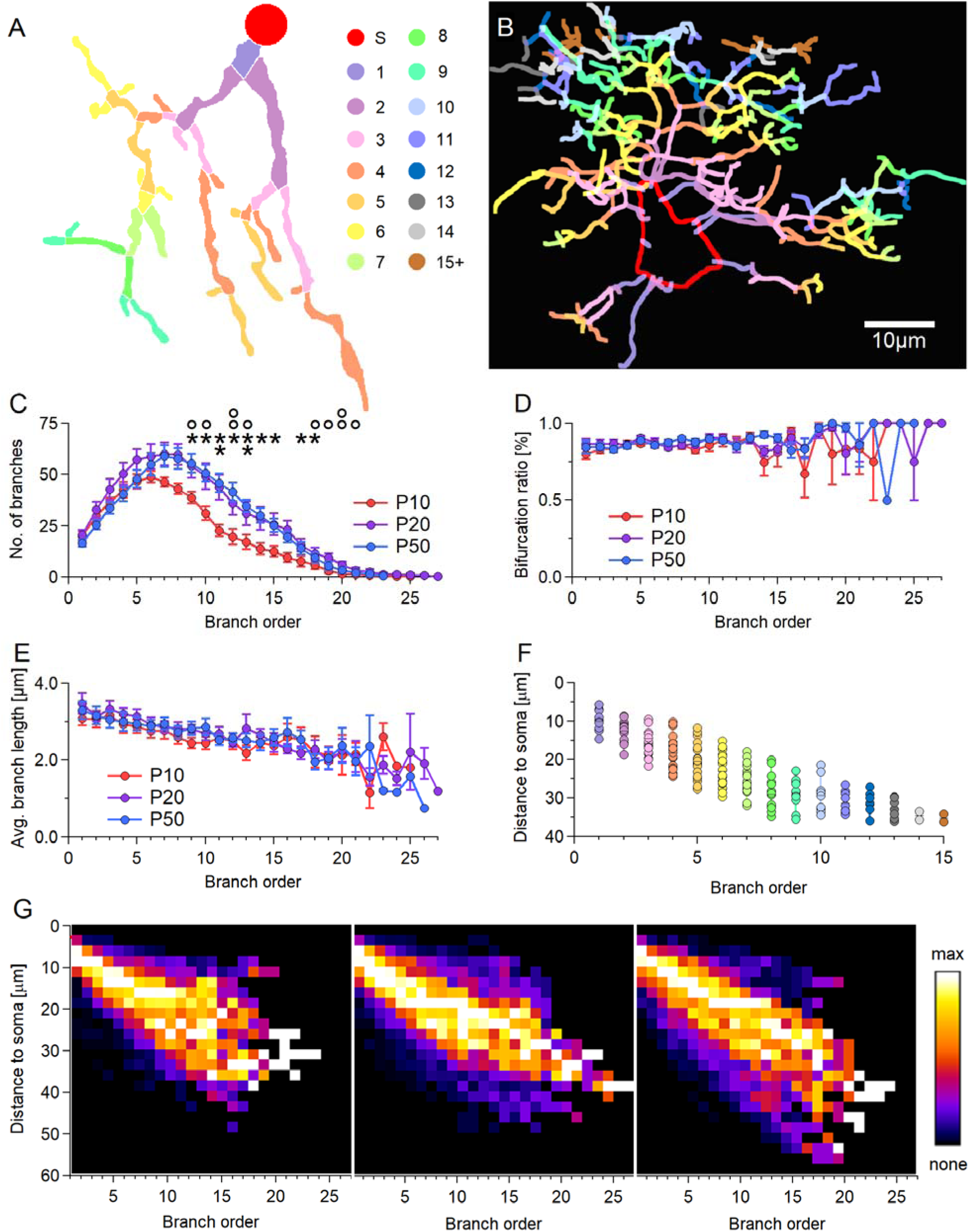
Larger number of higher-order branches underlies stronger branching of OPC processes in more mature animals. **(A):** Schematic representation of an OPC process with each branch colored in accordance to its position on the process (branch order). The color code used in this figure, figure (B) and (F) is the same and is shown to the right of the process. Soma is labeled in red. **(B):** Reconstruction of the whole OPC with each branch colored in accordance with its position on the process. **(C):** Comparison of the number of branches at each branch order. All points on the graph represent group averages ± SEM. Significant differences are marked with ° for P10 vs P20 comparisons; * for P10 vs P50 comparisons. °,* = p<0.05; °°,** = p<0.01. There were no significant differences between P20 and P50 groups. P10 group: n=13, N=7; P20 group: n=12, N=8; P50 group: n=13, N=8 animals. The same labeling, number of cells, and the number of animals apply to other graphs within this figure. **(D):** Comparison of the fraction of bifurcations at each branch position. **(E):** The average length of a single branch at each branch position. **(F):** Relationship between branch position on the process and the distance of the branch from the cell soma. Branches of the cell shown in (B) are separated by their order and plotted against distance between their origin point and the centroid of the soma. All branches of order 15 and higher are plotted together as 1 category. Black bar shows the median. **(G):** Comparison of the relationship between the position of a branch and its distance from the soma. At each order, the branch distance to soma is binned into 2.5 μm categories generating a histogram. Those histograms are plotted in an ascending order with the peak of the histogram colored white, zeroes colored black.

First, we analyzed the number of branches. We found that in the P10 group of animals, the number peaked (48.846 ± 2.396) at the branch order #6 after which the distribution declined (Figure 5C). The distributions in the P20 and P50 groups peaked at the branch order #7: 59.917 ± 5.610 at P20; and 59.077 ± 5.454 at P50 (Figure 5C). Afterwards, both distributions started to decline, at a similar rate, and became indistinguishable from each other (Figure 5C; p=0.067 to p=0.999). The statistically significant differences between the groups first appeared at the branch order #9, where the P10 group tested as different from both P20 and P50 (Figure 5C; 38.615 ± 2.688 at P10; 54.167 ± 5.934 at P20; 55.385 ± 4.460 at P50; P10 vs P20 p=0.0395; P10 vs P50 p=0.033; P20 vs P50 p=0.850).

Second, we sought to investigate possible differences between the groups in the branching pattern. We looked into the ratio of bifurcations among the branching points, plotting it against the branch order. We found that those ratios remained conserved throughout the branch orders and throughout the development (Figure 5D). For instance, at branch order #1 they were ∼0.85 in all groups (0.801 ± 0.034 at P10; 0.864 ± 0.035 at P20; 0.880 ± 0.023 at P50), and at branch order #13 they were ∼0.90 in all groups (0.913 ± 0.028 at P10; 0.893 ± 0.027 at P20; 0.904 ± 0.022 at P50). At branch orders above #13 the measurements become unreliable due to very low counts of branches. The distributions did not test as statistically significant at any point (p=0.057 to p=0.898).

Third, we investigated whether there are any distinguishable differences between the groups in the patterns of the average branch length. We found that the longest branches were located at the origin of the processes, i.e. closest to cell soma, and their length gradually decreased with each new branching order (Figure 5E). Thus, the length of the branches within the branch order #1 was on average ∼3.3 μm (3.081 ± 0.176 at P10; 3.467 ± 0.270 at P20; 3.300 ± 0.228), while the length of the branches within the branch order #13 was almost 30% shorter (2.182 ± 0.188 at P10; 2.471 ± 0.111 at P20; 2.504 ± 0.172 at P50). Moreover, the differences between groups were minor and not significantly different (p=0.104 to p=0.999; Holm-Šídák’s, Dunnett’s T3 or Dunn’s tests).

Finally, we combined the Branch Order Analysis and the Sholl analysis to visualize how the location of branches of various orders is related to their location in space. We measured the straight-line-distances between each branching point and the centroid of the cell soma, and plotted those distances vs the branch order. An example of such a plot for the cell in Figure 5B, is presented in Figure 5F, using the same color code for both panels. We found an almost linear relationship between the branch order and the distance from the cell soma (Figure 5G). In all groups, the branches of the higher order (#13+) were mainly located at more outer parts of the OPC cellular domain (Figure 5G). The distributions were also fanning out at higher branch orders pointing to higher variability of the data points.

Taken together, our findings demonstrate that larger cell domains and more extensive branching of OPC processes in mature animals is caused by an increase in the number of higher-order branches, rather than branch elongation.

### 3.6. Local branching density and neighbor distances are preserved across development

To assess whether increased branching in more mature OPCs is accompanied by changes in local branching density or spatial proximity between neighboring branching points, we performed nearest neighbor analyses (see Methods 2.9; Figure 6A-C). Two complementary approaches were used: first, quantification of the number of neighboring branching points within defined spatial cutoffs (1-5□μm), and second, measurement of Euclidean distances to the five nearest neighbors of each branching point. Results were analyzed relative to both the distance of the branching point from the soma centroid and its branch order.

**Figure 6:**
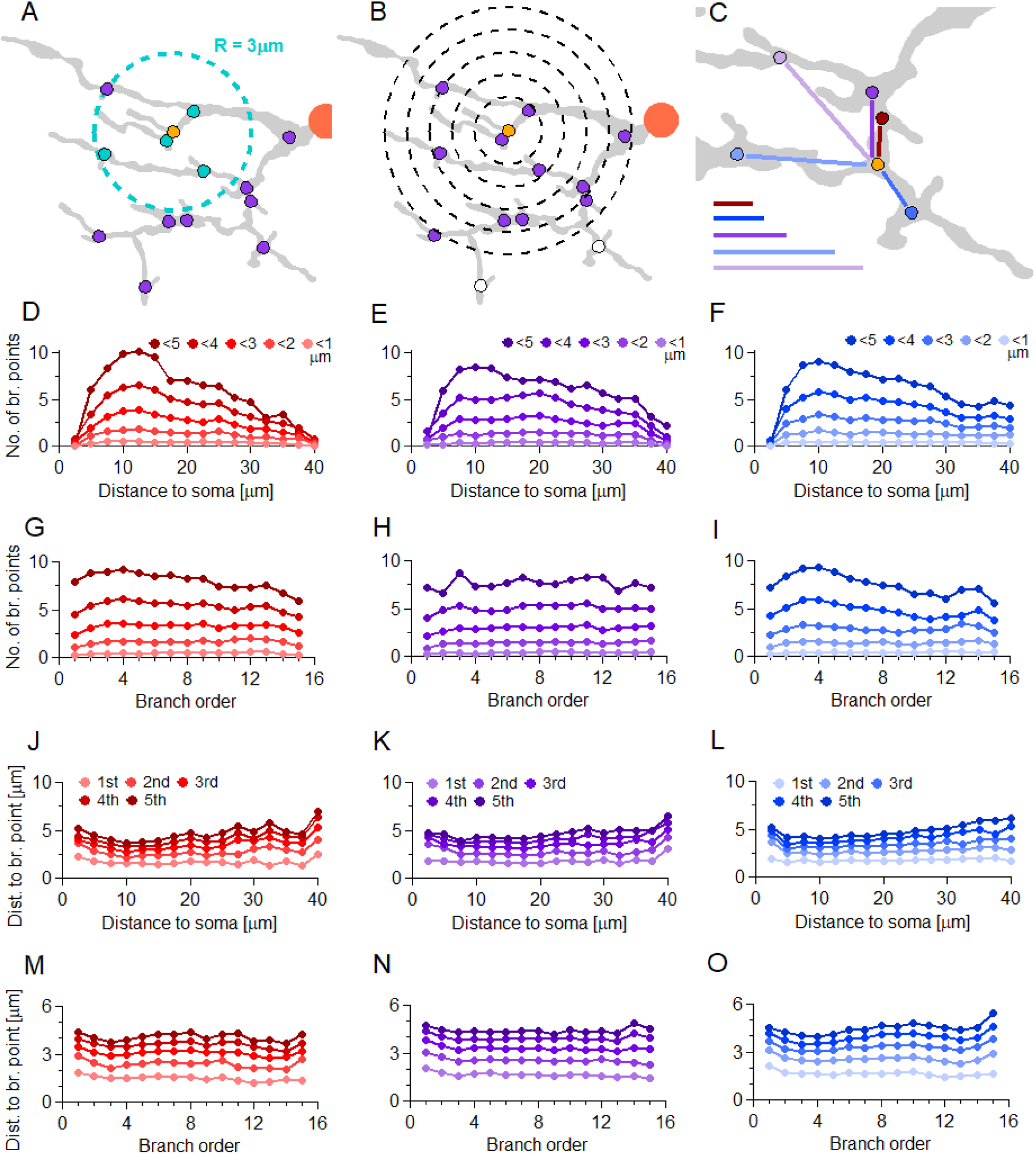
Number of and distances between neighboring branching points are developmentally preserved. **(A):** Schematic drawing of a single process with a selected branching point (orange), its closest neighbors within 3□μm (turquoise), and other branching points of the process in violet. The process origin at the soma is represented by a large red circle. **(B):** Principle of Sholl analysis for branching points. A selected branching point (orange) is placed at the center of five concentric shells, spaced 1□μm apart. All neighbors within 5□μm of the central point are shown in violet; all points beyond the largest shell are shown in white. **(C):** Distance to the five closest neighbors of the selected point (orange). Lines to the left represent the Euclidean distance between the selected point and each neighbor. **(D) :** Mean number of neighbors of a branching point within 0–5□μm, plotted against the distance of the point from the soma centroid (binned every 2.5□μm), as in (B), for the P10 age group. n=13 cells, N=7 animals. **(E):** Same as (D), for the P20 age group. n=12 cells, N=8 animals. **(F):** Same as (D), for the P50 age group. n=13 cells, N=8 animals. **(G):** Mean number of neighbors of a branching point within 0–5□μm, plotted against the branching order of the point, as in (B), for the P10 age group. n=13, N=7. **(H):** Same as (G), for the P20 age group. n=12, N=8. **(I):** Same as (G), for the P50 age group. n=13, N=8. **(J):** Euclidean distance between the selected point and its 1st to 5th closest neighbors, plotted against the distance of the point from the soma centroid (binned every 2.5□μm), as in (C), for the P10 age group. n=13, N=7. **(K):** Same as (J), for the P20 age group. n=12, N=8. **(L):** Same as (J), for the P50 age group. n=13, N=8. **(M):** Euclidean distance between the selected point and its 1st to 5th closest neighbors, plotted against the branching order of the point, as in (C), for the P10 age group. n=13, N=7. **(N):** Same as (M), for the P20 age group. n=12, N=8. **(O):** Same as (M), for the P50 age group. n=13, N=8.

In the first analysis, the number of neighboring branching points within 0-5□μm remained relatively stable across developmental stages when plotted against either the distance from the soma centroid (Figure 6D-F) or the branch order (Figure 6G-I). No consistent statistically significant differences were observed between P10, P20, and P50 groups at almost all distance bins, with the exception of 35-40□μm at the very end of the cell domain for P10 OPCs (Figure 6D-F; p□=□0.069 to p□=□0.999; 40□μm: p□=□0.0141 to p□=□0.0887; Holm-Šídák’s test) or by branch order (Figure 6G-I; p□=□0.082 to p□=□0.999, Holm-Šídák’s test), indicating that local neighbor density is maintained during development despite overall increases in process complexity.

Interestingly, in contrast to the previous analyses (Sections 3.4 and 3.5), the peak neighbor density fell around a distance of 10-12.5□μm from the soma (versus 15-17.5□μm; Figure 4C-F) and branch order #4 (versus order #6-7; Figure 5C), and remained highly consistent throughout the entire cell domain at cutoff diameters of 3□μm or lower (comparable to the average branch length of ∼2.78□μm; Figure 3H).

In the second analysis, the distances between each branching point and its first to fifth closest neighbors were plotted against the point’s distance from the soma (Figure 6J-L) and its branch order (Figure 6M-O). Again, no consistent significant differences were observed for either the branch order (p□=□0.051 to p□=□0.998; Holm-Šídák’s test) or the distance from the soma, and the few significant differences appeared largely incidental (p□=□0.208 to p□=□0.998; Holm-Šídák’s test). Mean Euclidean distances between neighboring branching points remained comparable across all groups, regardless of spatial location or hierarchical level within the process arbor.

These findings contrast with the results of the Sholl and Branch Order analyses (Sections 3.4 and 3.5), which showed significant developmental increases in process length, distal branching, and higher-order branch numbers. Together, the data suggest that although mature OPCs exhibit more extensive and distal arborization, the local spatial relationships between neighboring branching points - the immediate microarchitecture of the processes - are conserved. This indicates that the developmental increase in OPC complexity is driven by the addition of new, higher-order branches without altering the fundamental spacing or local density of branches.

### 3.7. The processes of OPCs show stronger preferential alignment with the lateral-medial than with the dorsal-ventral brain axis

During postnatal development and maturation of the animals, corpus callosum continues to grow, however, our measurements in the Allen and Paxinos brain atlases suggest that the growth is not uniform: corpus callosum seems to expand in the lateral-medial direction much more than in the dorsal-ventral direction. The volume gain may also not be uniform, and there is evidence for this in the human brain (Vannucci and others, 2017). Callosal growth likely influences the structural organization of OPCs. Furthermore, the myelination process within the corpus callosum that starts during the second postnatal week and continues into adulthood may also affect the morphology of OPCs and promote their processes to orient in the direction of the callosal axons. To test whether a preferential directionality of callosal OPCs’ processes exists, we investigated their alignment with the anatomical brain axes and compared it between the age-groups. For this analysis, we utilized a spherical coordinate system where the direction is given by two angles: a polar angle φ defined for x,y plane and an azimuth angle θ corresponding to z axis, perpendicular to both x and y (see Materials and Methods).

First, we investigated how many branches face the lateral-medial axis and dorsal-ventral axis (Figure 7A-C). The values were normalized to the total number of branches in each OPC, and averaged for a given age-group of animals. We found that in all three investigated groups, the large number of branches was closely aligned with the lateral-medial axis (0°-15°): 21.707 ± 1.753% for P10; 21.983 ± 0.985% for P20; and 27.056 ± 1.618% for P50 (Figure 7D-F). The P10 and P20 groups showed comparable distributions, both with significantly less branches than OPCs in the P50 group (Figure 7D-F; P10 vs P20 p=0.899; P10 vs P50 p=0.0472; P20 vs P50 p=0.0482, Holm-Šídák’s test). Notably, the more the branches misaligned with the lateral-medial axis the lower was their number, reaching a minimum number for branches aligned with the dorsal-ventral axis (75°-90°). OPCs in the P50 group had the lowest number of branches aligned with the dorsal-ventral axis, but the difference was statistically significant only between the P10 and the P50 group (Figure 7D-F, J; 14.370 ± 0.838% for P10; 13.463 ± 0.712% for P20; and 11.899 ± 0.454% for P50; P10 vs P20 p=0.360; P10 vs P50 p=0.0421; P0 vs P50 p=0.223, Holm-Šídák’s test).

**Figure 7:**
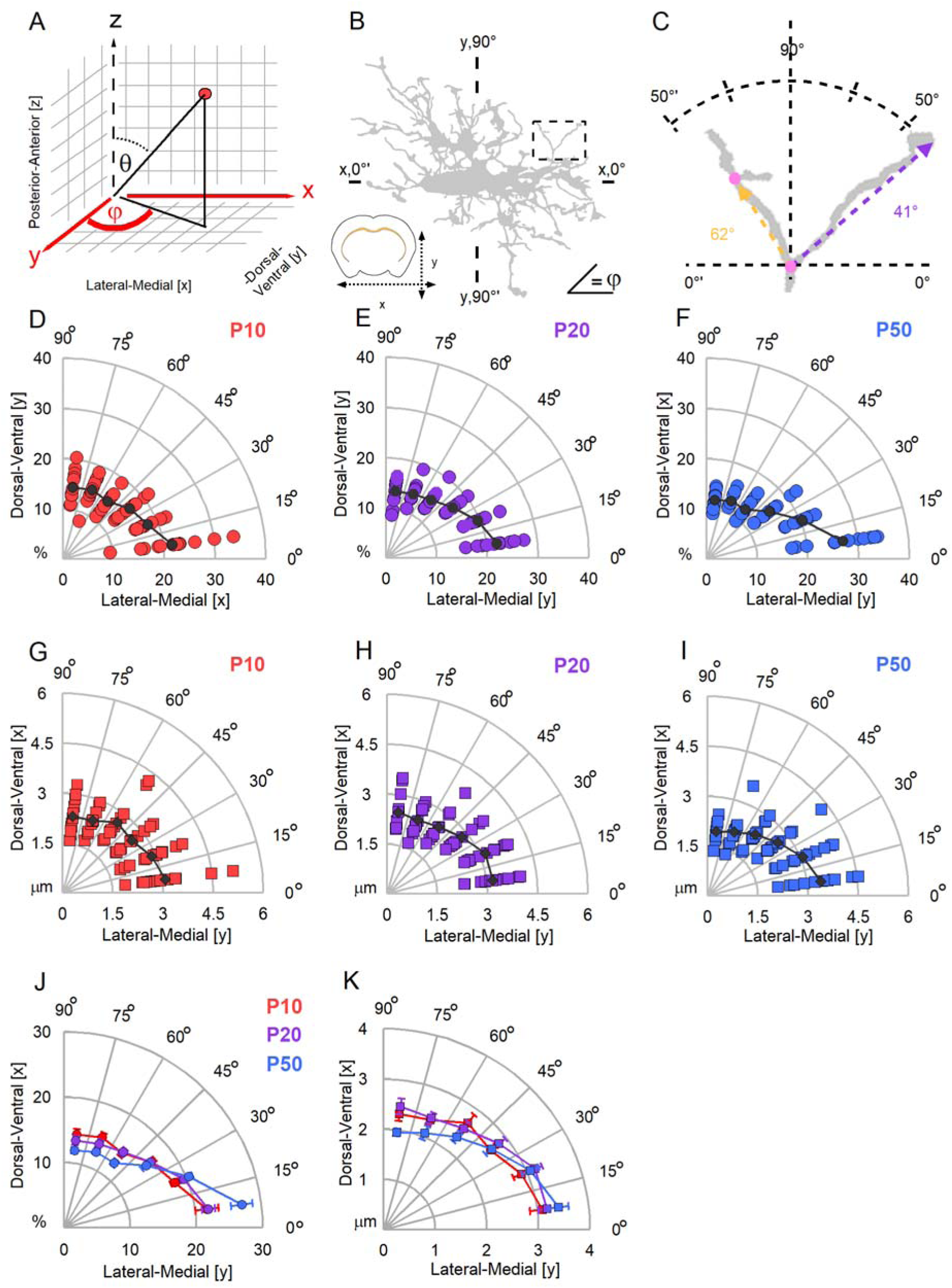
OPC processes preferentially align with and have longer branches when facing the medial-lateral over the dorsal-ventral body axis. **(A):** Medial-lateral and dorsal-ventral axes mapped onto a spherical coordinate system correspond to the x and y axes (colored red) and the planar angle φ between those axes. Note that the elevation angle θ is ignored in this analysis. **(B):** Example of a cell in the coronal view with angular values of φ added. Perfect alignment with the medial-lateral axis occurs when the value of φ=0°. Perfect alignment with the dorsal-ventral axis occurs when the value of φ=90°. **(C):** Inset from (B), showing enlarged part of the process with the φ values for 2 different branches. The direction of a branch is approximated by the direction of a vector spun between the beginning and the termination of the branch. Tortuosity of the branch is ignored. The color of the vector corresponds to the color of the φ measurement. **(D):** Distribution of branches in alignment with the medial-lateral or dorsal-ventral axes in the P10 age group. φ values are binned every 15° and plotted as a fraction (%) of all φ values measured per cell. Each red circle represents a percentage of φ values per 15° category in an individual cell. Black circle represents the group average, n=13, N=7 animals. **(E):** The same as (D) for the P20 age group, n=12, N=8 animals. **(F):** The same as (E) for the P50 age group. n=13, N=8 animals. **(G):** The average length of branches in alignment with medial-lateral or dorsal-ventral axes in the P10 age group. φ values are binned every 15° and branch lengths within each category are averaged. Each red circle represents the length of branches within a 15° category, measured in an individual cell. Black circle represents the group average, n=13, N=7 animals. **(H):** Similar to (G) for the P20 group, n=12, N=8 animals. **(I):** Similar to (G) for the P50 group, n=13, N=8 animals. **(J):** Summary of the proportion of branches in alignment with medial-lateral or dorsal-ventral axes in all age groups. Values for the P10 group are shown in red, for the P20 group in purple, and for the P50 group in blue. Each circle represents the group average ± SEM. Binning and fractions are the same as in (D)-(E). **(K):** Summary of the lengths of branches in alignment with medial-lateral or dorsal-ventral axes in all age groups. Values for the P10 group are shown in red, for the P20 group in purple, and for the P50 group in blue. Each circle represents the group average ± SEM. Binning and fractions are the same as in (G)-(I).

Next, we wondered whether there may be any correlation between the alignment of the branches along a given brain axis and their length. We found no statistically significant differences between the groups in the branch lengths, except for branches more closely aligned with the dorsal-ventral axis (60°-75°) where P50 cells had significantly shorter branches when compared with the P10 and P20 cells (Figure 7G-I, K; 2.360 ± 0.0959 μm for P10; 2.403 ±0.118 μm for P20; 1.961 ± 0.0786 μm for P50; P10 vs P20 p=0.761; P10 vs P50 p=0.0138; P20 vs P50 p=0.011, Holm-Šídák’s test). When we analyzed the ratios of lengths of the dorsal-ventral and the medial-lateral aligned branches, we found that they were nearly identical for P10 and P20 OPCs: 0.794 ± 0.069 for P10 and 0.791 ± 0.052 for P20 (p=0.999, Dunn’s test). However, for P50 OPCs the ratio was smaller (0.587 ± 0.026), and differed significantly from P10 and P20 cells (P10 vs P50 p=0.0219; P20 vs P50 p=0.0170, Dunn’s test).

### 3.8. OPC processes become more aligned with the posterior-anterior brain axis during postnatal development

To continue the analysis of the preferential orientation of the OPCs’ processes, we analyzed the azimuth angle θ (Figure 8A). In this analysis, the closer the θ angle of a branch is to 0°, the more that branch aligns with the posterior-anterior axis; the closer the θ angle of a branch is to 90°, the more that branch miss-aligns with the posterior-anterior axis (regardless of the x-y alignment) (Figure 8A-C).

**Figure 8:**
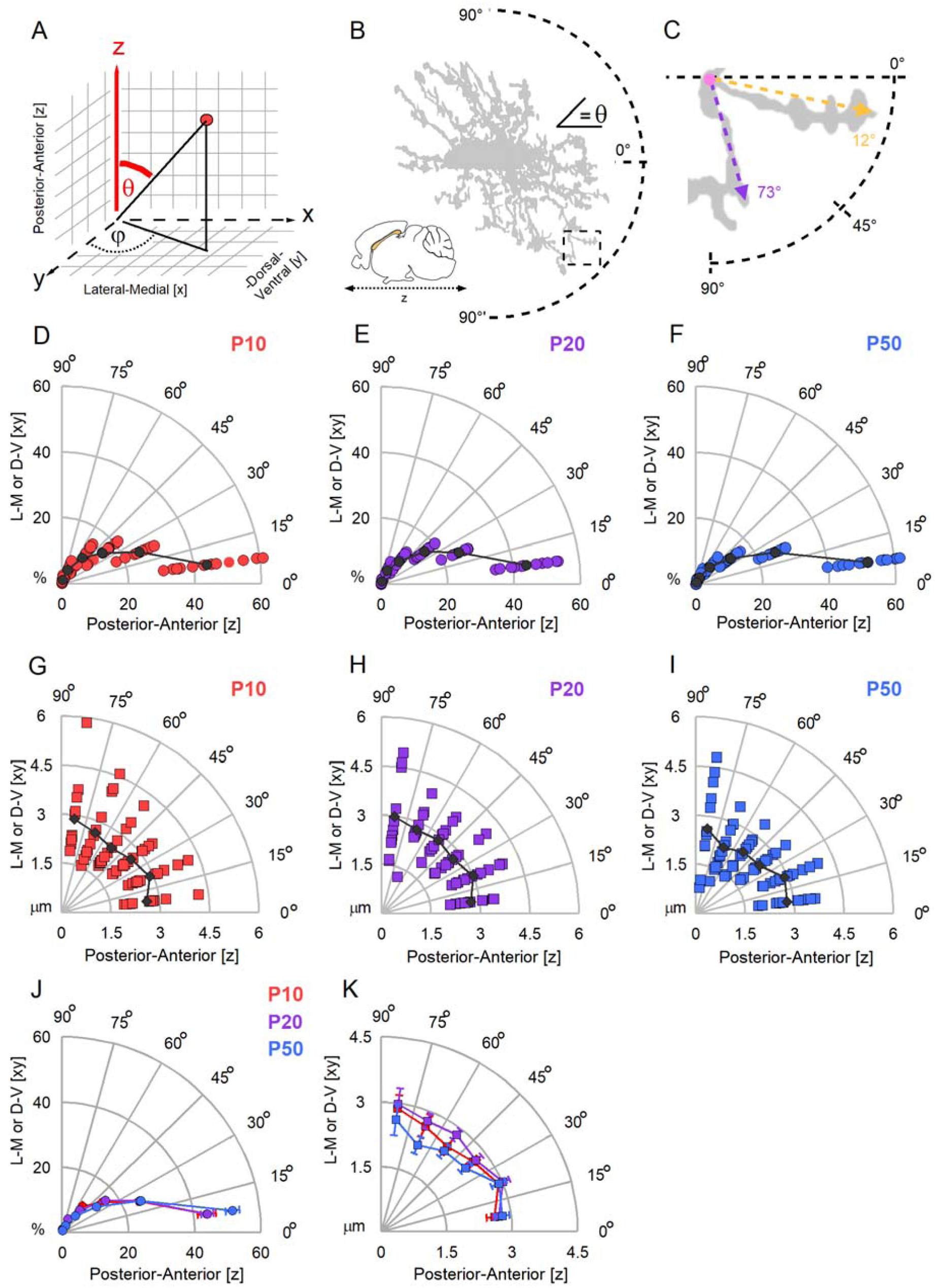
The processes of OPCs show stronger preferential alignment with the posterior-anterior brain axis than with the lateral-medial or dorsal-ventral brain axes. **(A):** Anterior-posterior axis mapped onto a spherical coordinate system corresponds to the z axis (colored red) and the elevation angle θ. Note that the planar angle φ is ignored in the analysis. **(B):** Example of a cell in a sagittal view with angular values of θ added. Perfect alignment with the anterior-posterior axis occurs when the value of θ=0°. The inset is enlarged in C. **(C):** Inset from (B) showing an enlarged part of the process with the θ values for 2 different branches. The direction of a branch is approximated by the direction of a vector spun between the beginning and the termination of the branch. Tortuosity of the branch is ignored. The color of the vector corresponds to the color of θ measurement. **(D):** Distribution of branches in alignment with anterior-posterior axis in the P10 age group. θ values are binned every 15° and plotted as a fraction (%) of all θ values measured per cell. Each red circle represents a % of θ values per 15° category in an individual cell. Black marker represents the group average, n=13, N=7. **(E):** Similar to (D) for the P20 age group, n=12, N=8. **(F):** Similar to (D, E) for the P50 age group, n=13, N=8. **(G):** The average length of branches in alignment with the anterior-posterior axis in the P10 age group. θ values are binned every 15° and branch lengths within each category are averaged. Each red circle represents the length of branches within a 15° category, measured in an individual cell. Black circle represents the group average. **(H):** Similar to (G) for the P20 age group, n=12, N=8. **(I):** Similar to (G) for the P50 group, n=13, N=8. **(J):** Summary of the proportion of branches in alignment with anterior-posterior axis in all age groups. Values for the P10 group are shown in red, for the P20 group in purple, and for the P50 group in blue. Each circle represents the group average ± SEM. Binning and fractions are the same as in (D)-(E). **(K):** Summary of the lengths of branches in alignment with anterior-posterior axis in all age groups. Values for the P10 group are shown in red, for the P20 group in purple, and for the P50 group in blue. Each circle represents the group average ± SEM. Binning and fractions are the same as in (G)-(I).

We found that high number of branches of OPCs’ processes align with the anterior-posterior axis in all groups, but P50 OPCs had more anterior-posterior aligned branches than P10 or P20 cells, and the differences were statistically significant: 43.913 ± 2.787 for P10; 44.070 ± 1.863 for P20; 51.774 ± 2.159 for P50; (Figure 8D-F; P10 vs P20 p=0.761; P10 vs P50 p=0.0138; P20 vs p50 p=0.0110, Holm-Šídák’s test). However, when we investigated the average lengths of the branches we found no significant differences between the groups (Figure 8G-I, K; P10 vs P20 p=0.458 to 0.978; P10 vs P50, p=0.296 to 0.971, P20 vs P50 p=0.160 to 0.953, Holm-Šídák’s test).

### 3.9. The geometry of successive branches in OPCs’ processes: most branches change direction

Dendrites of neurons branch extensively, and often the elaboration of branches is not random but occurs at a preferential angle (Cherniak, 1992; Leguey and others, 2016; Rojo and others, 2016). Hence, we were wondering whether OPCs elaborate branches at specific angles, and whether the angles are the same throughout the whole process arbor. To address this question, we studied changes in the direction between the successive branches by measuring the planar angle between pairs of branches which originate from the same branching point (Figure 9A, B). In the analysis, 0° indicates no change in the direction while 180° indicates complete reversion of the direction.

**Figure 9:**
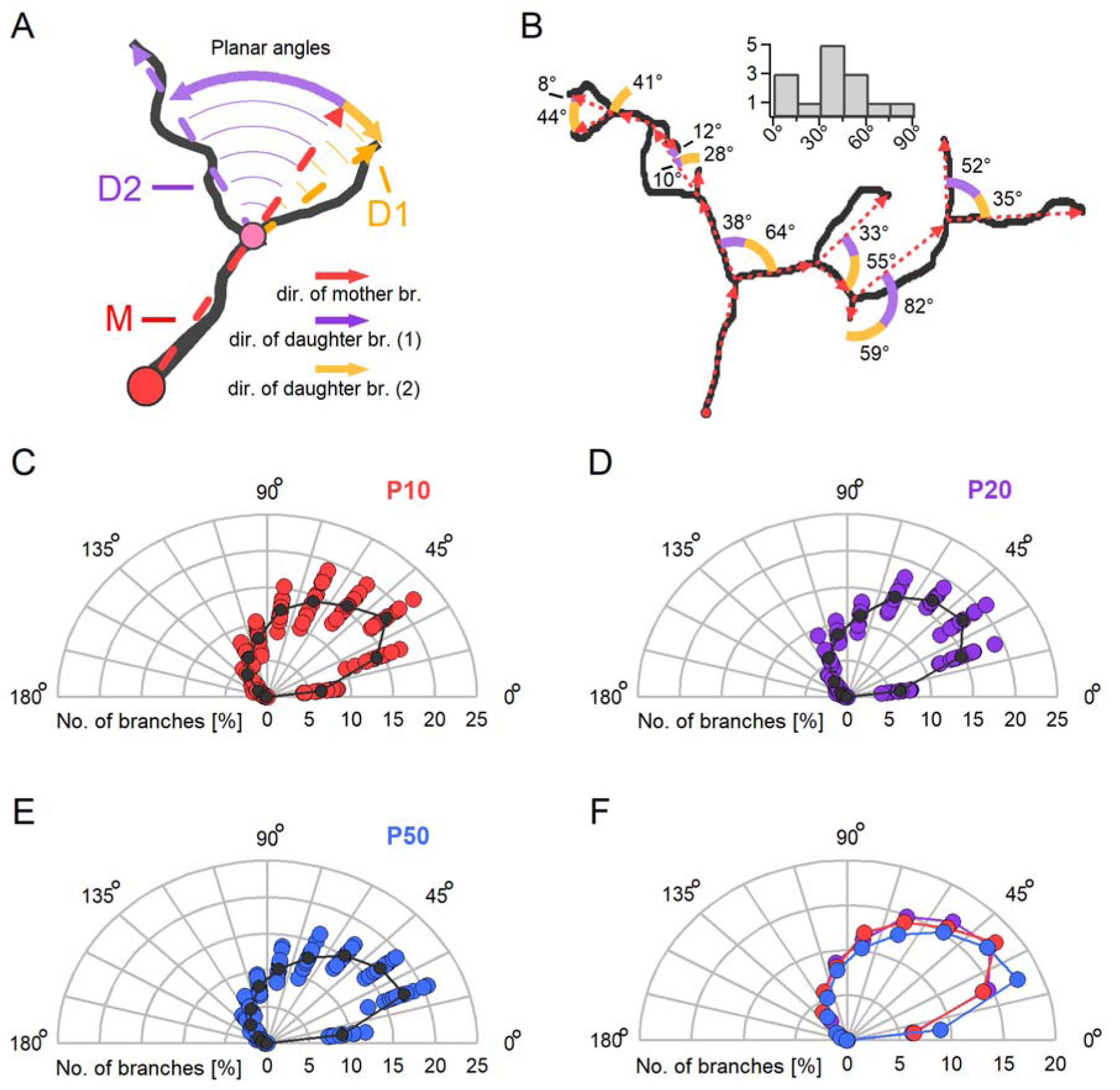
The majority of new branches show a direction preference, which is preserved during animal maturation. (A): Example of planar angle measurement to estimate the change in branch direction. The change of direction is measured at a branching point (pink circle) as an angle between the direction vector of the mother branch (M, red) and the direction vectors of its daughter branches (D1, orange; D2, violet). The angle measurements are indicated by colored arc arrows, color corresponding to respective daughter branches. **(B):** Example of a complex process with the changes to branch direction measured for every branch. Similar to (A), angular measurements for the first daughter branch are colored orange and for the complementary daughter branch in violet. Direction vectors for each branch are in red. The planar angle measurements for the example, binned every 15°, from 0° to 90°, are summarized in a histogram above the graph. **(C):** Distribution of the planar angle values measured in the P10 age group. Measurements are binned every 15°, with 0° indicating no change in branch direction and 180° indicates complete reversion in direction. Each red circle represents a % of angular values per 15° category in an individual cell. Black circle represents the group average, n=13, N=8 **(D):** Similar to (C) for the P20 age group, n=12, N=8. **(E):** Similar to (C) for the P50 age group, n=13, N=8. **(F):** Summary of (C)-(E) with group averages plotted together.

We found that in all three experimental groups, the majority of branches changed direction by a significant degree. In the P10 and P20 groups, about 16% of branches (15.9 ± 0.515% for P10; 16.508 ± 0.300% for P20) change the direction by 45°-60°, about 17% of branches (17.765 ± 0.535% for P10; 17.047 ± 0.538% for P20) change the direction by 30°-45°, about 14% of branches (14.096 ± 0.533% for P10; 14.678 ± 0.576% for P20) changed the direction by 15°-30°, and only about 6% of branches (6.378 ± 0.392% for P10; 6.186 ± 0.312% for P20) change the direction by less than 15° (Figure 9C-F). There was no statistically significant difference in those values between the P10 and P20 cells (Figure 9C-F; 0°-120°, one-way ANOVA, p=0.268 to 0.674).

In the P50 group, more branches appear to elaborate at shallower angles than in P10 and P20 groups: ∼17.164 ± 0.393% of branches change the direction by 30°-45°, another 17.283 ± 0.649% of branches change the direction by 15°-30°, and 8.834 ± 0.378% branches changed the direction by less than 15° (Figure 9C-F). The difference between P10 and P50 groups was statistically significant for 0°-30° (p=0.0000554 to 0.00133, Holm-Šídák’s test), while the difference between P20 and P50 groups was statistically significant till 75°(p=0.0000365 to 0.0368, Holm-Šídák’s test).

Finally, we tested whether changes of the branch direction are related to the branch order. For this, we measured the planar angle for all branches of each successive order (Figure 10A). We found almost no significant differences in the angle medians, neither within nor between groups (Figure 10B-D; p=0.426 to p=0.999, one-way ANOVA), with a single significant difference at branch order #13 (maximal br. order for some P10 cells; P10 vs P20 p=0.0145; P10 vs P50, p=0.0145; Holm-Šídák’s test) suggesting that OPCs preserve their branching patterns irrespective of the branch order. Interestingly, we noted a minor increase in the planar angles with successive orders in the P10 and P20 groups (although not statistically significant), but not in the P50 group.

**Figure 10:**
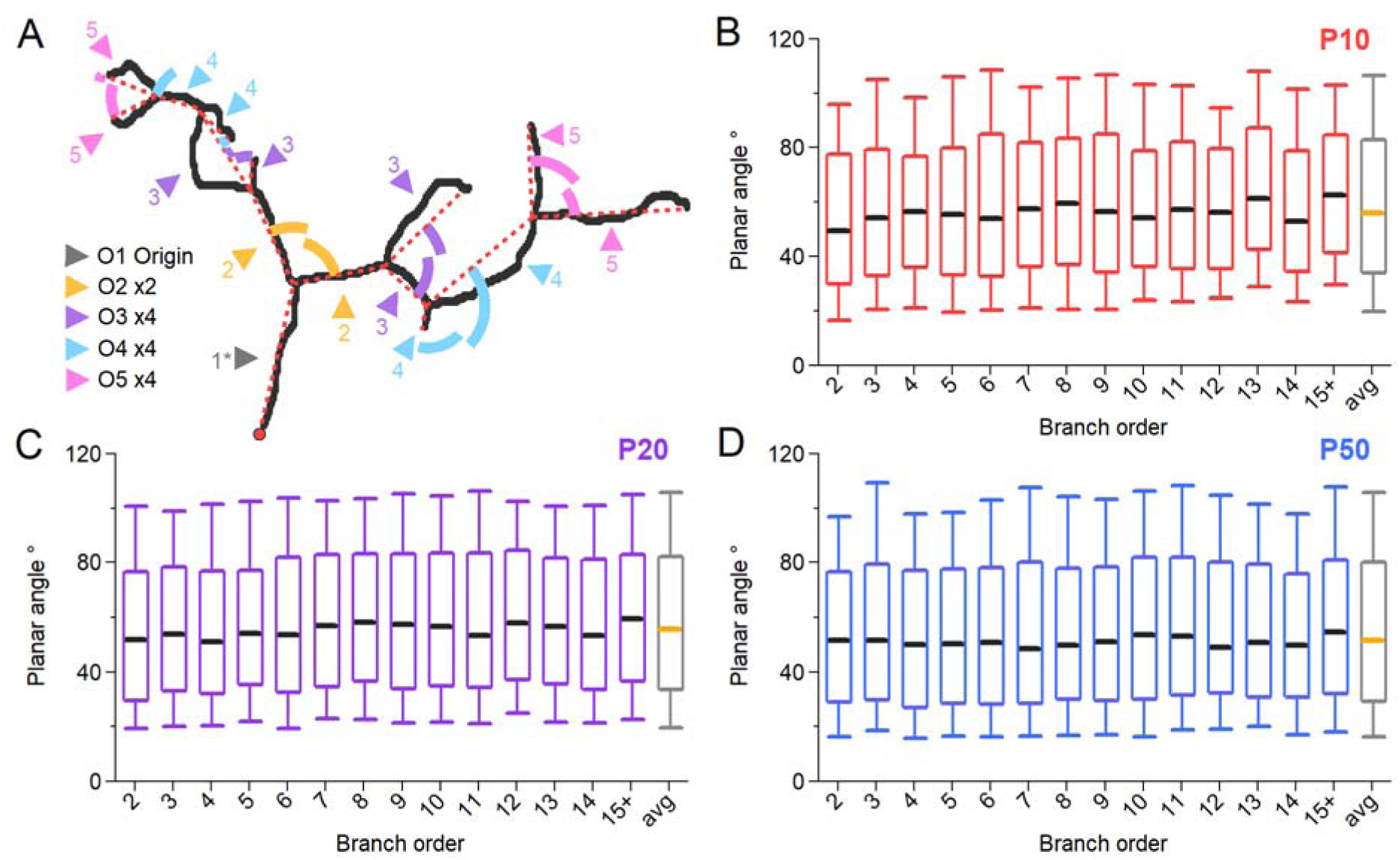
Changes in branch direction are not dependent on the position within a process (branch order) and do not change during development. **(A):** Example of a complex process of an OPC with the changes to branch direction measured for every branch color coded by branch order: order 1 in gray, order 2 in orange, order 3 in violet, order 4 in turquoise, order 5 in pink. Arrowheads point towards the branch for which the angle is measured and are colored based on the branch order. **(B):** Box plots summarizing the change in direction at increasing branch orders in the P10 age group. Each red box represents the 25^th^ to 75^th^ percentile, whiskers represent the 10^th^ and 90^th^ percentile, and the black bar represents the median. Grey box plot represents the average among all orders, with the median in orange, n=13, N=7. **(C):** Similar to (B) for the P20 age group, n=12, N=8. **(D):** Similar to (B, C) for the P50 age group, n=13, N=8.

## 4. Discussion

In the present study, we used transgenic mice expressing *membrane-tagged* GFP in OPCs, high-resolution imaging, and detailed quantitative morphometric analysis to investigate the geometrical principles that govern structural organization of OPCs in the mouse corpus callosum. Our major findings are: (1) During the first two months of postnatal life in mice, the total length of all OPCs’ processes increases; the increase is caused by elaboration of additional branches from the existing processes rather than by the appearance of new processes; (2) New branches are preferentially added to more distal sites of OPCs’ processes and, as a result, an increase in arborization of OPC processes in adult animals is caused by larger number of higher-order branches; (3) The processes of OPCs show stronger preferential alignment with the posterior-anterior brain axis than with the lateral-medial or dorsal-ventral brain axes; at the same time, the processes of OPCs show stronger preferential alignment with the lateral-medial than with the dorsal-ventral brain axis.

### 4.1. During animal maturation, the total length of white matter OPC processes increases, primarily through addition of new branches to the existing processes

The first key finding of our study is that the total length of callosal OPC processes increases during the first two months of postnatal life in mice, primarily through the elaboration of additional branches from existing processes, rather than the formation of entirely new trees of processes.

Why do OPCs increase the total length of their processes? An increased complexity of the OPCs’ arbor may reflect a shift in their functions and/or behavior, and/or appearance of new functions in the juvenile and adult brain in addition to those which OPCs already have in a very young brain. We observed that the total length of processes was larger in P20 and P50 groups compared to the P10 group. During the third postnatal week in mice, an extensive myelination wave takes place in the corpus callosum (Chen and others, 2018; Peris and others, 2023; Son and others, 2017); at the age of two months, myelination continues although at a much slower rate (Sturrock, 1980; Tripathi and others, 2017). Hence, refinement and elongation of the OPCs’ arbors in the P20 and P50 age-groups may reflect the search of OPCs for new axons as myelination targets, or may be necessary to ensure that sufficient number of processes is available for differentiation and elaboration of new myelin sheaths. Besides, less complex trees of processes in OPCs of the young animals (P10 group) may reflect their higher division rate, in line with our previous study which suggested that OPCs retract many processes during metaphase of mitosis and re-elaborate the processes in telophase (Kukley and others, 2008). It is also possible that during early postnatal development, OPCs focus on expansion and migration, while later during postnatal maturation the environmental monitoring is emerging as their more important function. This idea is also consistent with a previous observation *in vivo* showing that processes of mature OPCs retain dynamic behavior (Hughes and others, 2013).

But why do OPCs elaborate new branches from the existing processes rather than build the entire new trees of processes? A progressive refinement of OPCs’ arbors rather than a wholesale restructuring of their morphology may have several reasons. First, based on the information about neuronal dendrites and axons, extending new branches is likely structurally more efficient and require lower energy costs because the existing processes already contain all the necessary cytoskeletal and molecular infrastructure including actin, microtubules, membrane trafficking machinery, and organelles (Arikkath, 2012; Curran and others, 2024; Dent and others, 2011). Cell surface receptors, cell adhesion molecules, signaling molecules, secreted factors, and ion channels, e.g. receptor tyrosine kinases and GPSRs, which mediate the effects of the extracellular signals such as e.g. neurotransmitters and growth factors required for building new branches, may also be already present on the membrane of the existing processes (Arikkath, 2012). Second, branching from existing processes likely allows the OPCs to respond quickly to multiple local cues, e.g. growth factors, axonal activity (Arikkath, 2012; Hamad and others, 2023; Shah and others, 2010). Growing entirely new trees of processes may take much longer to initiate, and may require re-orientation of cell polarity, activation of cell soma signaling at multiple levels, etc. Finally, there may be spatial constraints: creating a new tree of processes may lead to a larger unnecessary overlap of several processes, while elaborating multiple new branches is likely to result in a finer targeted exploration of the local extracellular space. Interestingly, dendritic and axonal arbors of neurons in various species are capable of self-avoidance to prevent overlap of processes, and arbors of neurons that innervate the same area are often arranged in a tiled pattern that maximizes coverage of that area while minimizing overlap between neighboring arbors (Grueber and Sagasti, 2010; Jan and Jan, 2010). Tiled patterns have also been described for the OPCs and astrocytes, although the mechanisms of tiling remain less investigated in glial cells than in neurons (Barber and others, 2021).

Does it mean that OPCs establish their arbors once and never build the entirely new trees of processes at all? Probably it is not the case. But it seems likely that elaboration of the new trees of processes by OPCs does not occur as a function of age, but takes place during other events which OPCs may undergo. For example, OPCs elaborate new trees of processes after mitosis which they may undergo both in the developing and also in the adult brain (Kukley and others, 2008; Marisca and others, 2020). It is also possible that OPCs may remodel their whole arbor of processes and add new trees during differentiation and after injury (Hughes and others, 2013; Li and Leung, 2015; Marisca and others, 2020).

### 4.2. New branches preferentially appear at distal sites of the processes of OPCs

The second key observation of our study is that in the OPCs of the more mature animals, new branches preferentially appear at distal sites of the processes, resulting in a larger proportion of higher-order branches in the arbors of adult OPCs. There may be several reasons why OPCs are more likely to elaborate new branches at the distal sites of their processes rather than in the proximity to the cell soma. First, there may be greater cytoskeletal plasticity at the distal sites, e.g. higher actin turnover and microtubule dynamics, while proximal regions may be more structurally stabilized and less responsive to various remodeling signals. Second, targeted membrane and organelle trafficking to the distal sites of the processes may ensure better availability of the membrane and protein components for elaboration of the new branches. In addition, mitochondria and endoplasmic reticulum present within the processes of OPCs may be efficient in supporting energy demands and local synthesis during extension of the new branches, in analogy to their role for branching in neurons (Lanoue and Cooper, 2019; Winkle and others, 2016). For instance, in neuronal cultures branching of sensory axons has been shown to occur at sites populated by stalled mitochondria, and the endoplasmic reticulum co-localizes with mitochondria at sites of branching (Spillane and others, 2013). Large proportion of axon branching sites exhibited mitochondria also in an acute living embryonic spinal cord explant preparation (Spillane and others, 2013). Mitochondria and endoplasmic reticulum are present also in neuronal dendrites and play a role in dendritic branching (Lanoue and Cooper, 2019; Virga and others, 2024). Mitochondria in the processes of OPCs are mobile, and their preferential localization changes when OPCs differentiate into myelinating oligodendrocytes (Bame and Hill, 2024), but their role in OPCs branching remains to be investigated. Functionally, elaboration of new branches at the distal sites of the arbor rather than in the proximity to the cell soma may be necessary to extend the reach of OPCs into the new areas of brain parenchyma, and to expand the volume for sensing the extracellular signals and responding to them. It may also have an impact on more efficient sampling of neuronal activity, particularly during activity-dependent myelination. Notably, the mode of arbor growth by elaboration of additional branches from the distal sites also parallels the dendritic maturation observed in certain neuronal types, where distal elaboration enhances functional compartmentalization and information integration (Jan and Jan, 2010).

### 4.3. The processes of callosal OPCs preferential align with the posterior-anterior brain axis and to a lesser extent with the lateral-medial axis

The third key finding of our study is that branches of OPC processes in the corpus callosum exhibit a non-random, preferential alignment with the posterior-anterior brain axis, and to a lesser extent with the lateral-medial axis over the dorsal-ventral axis.

The corpus callosum is composed of the commissural axons of long-range projection neurons that travel from one hemisphere to the other, i.e. along the lateral-medial axis. Hence, it appears logical that callosal OPCs align their processes in the direction of the lateral-medial axis, i.e. parallel to the callosal axons. This orientation is most likely supported by mechanical and biochemical cues. For instance, the contact guidance may promote the cytoskeletal alignment of the OPCs’ processes along the dominant course of the fibers, and this orientation likely gets reinforced by physical alignment of the extracellular matrix molecules and axonal membranes. It is possible, that this directionality of the OPCs processes is set up already during early development when axon guidance molecules (e.g. ephrins, netrins) establish the midline-crossing direction of the callosal axons (Fame and others, 2011; Nishikimi and others, 2013; Silver and others, 1982), and OPCs migrating to the corpus callosum (Kessaris and others, 2006; Tsai and others, 2016) are exposed to the same/similar molecular cues and molecular gradients. Functionally, alignment of the processes of OPCs with the course of callosal axons may maximize the contact between the axons and the OPCs for more efficient axon-OPCs synaptic signaling, for search and stabilization of the future myelination targets by OPCs.

Interestingly, however, our study shows that, although OPCs align branches of their processes with the lateral-medial axis more readily than with dorsal-ventral axis, the lateral-medial orientation is not their most preferred orientation. We found that a much higher proportion of branches in OPCs’ processes preferentially align with the posterior-anterior brain axis. There may be several explanations for this finding. Although callosal axons themselves run mostly parallel to the lateral-medial axis, the corpus callosum as an anatomical brain structure extends in the anterior-posterior direction (Wang and others, 2020). Therefore, positioning their branches along the anterior-posterior axis may allow OPCs to cross over multiple axons touching them or establishing contacts with them, thus sampling axonal signals and different layers of the callosal bundle. These OPC-axon contacts may be brief, transient events (Hughes and others, 2013; Kirby and others, 2006), helping a given OPC to explore as many axons as possible sensing axonal diameter, neuronal activity, readiness for myelination, etc. Alternatively, these axon-OPCs contacts may be more stable connections reflecting the establishment of synaptic signaling between axons and OPCs. If any single OPC was to receive synaptic input from axons throughout several layers of the callosal bundle rather than from the axons within the same layer, this would allow each OPC to act as a more comprehensive reader and integrator of the neuronal activity, probably resulting in a broader axon-glia network activity and higher level of synchronization of axon-glia events. This may be beneficial for energy optimization, and may allow for more efficient synchronization of proliferation and differentiation of OPCs by axons (Karadottir and Kuo, 2018; Nagy and others, 2017). Orientation of OPCs’ branches in anterior-posterior direction may be beneficial for a well-organized filling of the 3D volume of the white matter and allow to avoid overlapping with neighboring OPCs, while at the same time ensuring uniform surveillance and coverage of the axonal targets by OPCs (Barber and others, 2021).

## 5. Conclusions

Taken together, our study supports a model in which the morphology of OPCs becomes increasingly complex and spatially refined with maturation of animals, probably enhancing the ability of OPCs to integrate into the mature CNS microenvironment. The observed structural changes are likely critical for supporting the expanded repertoire of OPCs functions in adulthood, including axon-glia synaptic signaling, homeostatic surveillance, and participation in repair mechanisms following injury.

Future studies should explore the functional significance and the molecular mechanisms driving the preferential elaboration of the distal branches, and the favored axis-specific orientation of the branches within the OPC arbor of processes. For instance, molecules playing a role in cytoskeletal organization of OPCs, growth factors, and components of extracellular matrix may play a role in regulation of the spatial orientation of OPCs and refinement of their arbors of processes. In future research, it would be also interesting to perform a detailed characterization of the morphological changes that occur in OPCs during pathological conditions, e.g. during demyelinating and re-myelinating events in various diseases. The goal of this research would be to find out how pathological changes in OPCs’ morphology differ from those during development and maturation, and whether therapeutic interventions aimed at modulating the morphology of OPC during diseases may enhance regeneration.

## Contributions

Conceptualization: B.K., M.K.

Data curation: B.K., M.K.

Formal analysis: B.K.

Funding acquisition: M.K.

Investigation: B.K.

Methodology: B.K.

Project administration: M.K.

Resources: M.K Software: B.K.

Supervision: M.K.

Validation: B.K., M.K.

Visualization: B.K.

Writing - original draft: B.K.

Writing - review & editing: B.K., M.K.

## Acknowledgements

We thank Dr. Bill Stallcup (Burnham Institute, USA) for the gift of NG2 antibodies. We thank Dr. Da Guo (Okinawa Institute of Science and Technology, Japan) and Dr. Botond Antal (University of Minnesota, USA) for their help with the nearest neighbor analysis. We thank Ms. Victoria Wedler for her help with immunohistochemistry, confocal imaging and Neurolucida tracings.

